# β-Cyclodextrins as affordable antivirals to treat coronavirus infection

**DOI:** 10.1101/2022.11.16.516726

**Authors:** Dalia Raïch-Regué, Raquel Tenorio, Isabel Fernández de Castro, Daniel Perez -Zsolt, Jordana Muñoz-Basagoiti, Martin Sachse, Sara Y. Fernández-Sánchez, Marçal Gallemí, Paula Ortega-González, Alberto Fernández-Oliva, José A. Gabaldón, Estrella Nuñez-Delicado, Josefina Casas, Ferran Tarrés, Júlia-Vergara Alert, Joaquim Segalés, Jorge Carillo, Julià Blanco, Bonaventura Clotet Sala, José P. Cerón-Carrasco, Nuria Izquierdo-Useros, Cristina Risco

## Abstract

The SARS-CoV-2 pandemic made evident that we count with few coronavirus-fighting drugs. Here we aimed to identify a cost-effective antiviral with broad spectrum activity and high safety and tolerability profiles. We began elaborating a list of 116 drugs previously used to treat other pathologies or characterized in pre-clinical studies with potential to treat coronavirus infections. We next employed molecular modelling tools to rank the 44 most promising inhibitors and tested their efficacy as antivirals against a panel of α and β coronavirus, e.g., the HCoV-229E and SARS-CoV-2 viruses. Four drugs, OSW-1, U18666A, hydroxypropyl-β-cyclodextrin (HβCD) and phytol, showed antiviral activity against both HCoV-229E (in MRC5 cells) and SARS-CoV-2 (in Vero E6 cells). The mechanism of action of these compounds was studied by transmission electron microscopy (TEM) and by testing their capacity to inhibit the entry of SARS-CoV-2 pseudoviruses in ACE2-expressing HEK-293T cells. The entry was inhibited by HβCD and U18666A, yet only HβCD could inhibit SARS-CoV-2 replication in the pulmonary cells Calu-3. With these results and given that cyclodextrins are widely used for drug encapsulation and can be safely administered to humans, we further tested 6 native and modified cyclodextrins, which confirmed β-cyclodextrins as the most potent inhibitors of SARS-CoV-2 replication in Calu-3 cells. All accumulated data points to β-cyclodextrins as promising candidates to be used in the therapeutic treatments for SARS-CoV-2 and possibly other respiratory viruses.

## Introduction

Almost three years have passed since the SARS-CoV-2 pandemic began. Novel vaccines developed against this particular coronavirus have changed the global landscape protecting from severe coronavirus disease (COVID-19) and decreasing the death toll faced at the early stages of the pandemic. Yet, it has taken more than a year for COVID-19 vaccines to reach 20% of the population in low-income nations. Even in wealthy countries, where 20% vaccination was achieved last April 2021, there is only about an 80% of vaccination rate nowadays (https://ourworldindata.org/covid-vaccinations). After vaccination, vulnerable and immunocompromised individuals are still at high risk upon infection (Benet et al., 2022). Unfortunately, all vaccines approved so far have failed to confer sterilizing immunity, and may decrease but not fully protect from infection.

Under this scenario, therapeutic approaches are still needed to protect those at higher risk after SARS-CoV-2 infection, which include individuals who lack access to vaccine regimens or those who are vaccinated but fail to mount adequate immune responses. Safe, affordable and effective antivirals could be of great help to mitigate these contingencies, offering treatment for those individuals not only at higher risk of developing severe COVID-19, but also for those with moderate or mild outcomes that could also benefit from reducing the time and disease-associated symptoms. WHO recommends different treatments that limit severe disease or risk of hospitalization (Agarwal et al., 2020). Some of these antiviral treatments are based on neutralizing antibodies or intravenously administered drugs that can be dispensed in the hospitals only. Therapeutic antibodies such as casirivimab-imdevimab have shown activity in clinical trials before the surge of new variants. However, pre-clinical studies suggest that this combination lacks neutralization activity against omicron (Agarwal et al., 2020). In the case of sotrovimab, although activity might be retained, higher concentrations of the antibodies would be needed for neutralization of omicron (Agarwal et al., 2020). Remdesivir targets the viral RNA polymerase but still requires intravenous treatment, although new orally-available formulations are about to be tested (Schäfer et al., 2022). Molnupiravir, another inhibitor of the viral RNA polymerase, is administered orally and like remdesivir, it has shown clinical benefits when administered early upon infection (Jayk Bernal et al., 2022). WHO weakly recommends administration of Molnupiravir, and only for patients at high risk of being admitted to hospital with COVID-19 (Agarwal et al., 2020). Paxlovid is taken orally, but is not widely accessible and imply costs that exceed those of vaccines (Buxeraud et al., 2022). Moreover, Paxlovid is only available for patients that tolerate Ritonavir as part of the treatment with the antiviral nirmatrelvir, and has also raised concerns as monotherapy due to associated viral resistance (Yang et al., 2022), which may limit efficacy against new variants of concern (VOC).

None of the antivirals currently approved have the cost-effective profile or the easy administration route needed to offer prophylaxis to those vulnerable individuals at higher risk of developing severe disease upon SARS-CoV-2 infection on a global scale. Nevertheless, the key to control the SARS-CoV-2 pandemic might follow an approach similar to HIV-1 Pre-Exposure Prophylaxis (PrEP), which has been employed to decrease the HIV-1 infection rates (Molina et al., 2022). With the success of this prior HIV-1 strategy, having an affordable antiviral with prophylactic potential and a broad-spectrum activity at the initial surges of novel VOC could be key to decrease SARS-CoV-2 transmission rates.

Here we aimed to identify a cost-effective antiviral with broad spectrum activity and high safety and tolerability profiles. We began elaborating a list of drugs previously used to treat other pathologies or characterized in preclinical studies with potential to treat coronavirus infections. We next employed molecular modelling tools to rank the most promising inhibitors and tested their efficacy as antivirals against two representative viruses from the α and β coronavirus genera: the HCoV-229E and SARS-CoV-2 viruses. With a combination of computational chemistry, virology, cell biology and electron microscopy methods, we studied 44 compounds. Four of them showed antiviral activity against both HCoV-229E and SARS-CoV-2, being β-cyclodextrins the most promising candidates to treat the infection caused by SARS-CoV-2.

## Material and methods

### Molecular modelling

#### Viral targets

The characterization of the crystal structure of the M^pro^ (Jin et al., 2020), an enzyme recruited by the SARS-CoV-2 to complete the replication and transcription steps, was included in the protein data bank (PDB) library (code 6LU7) and allowed the use of virtual screening (VS) techniques (Berman et al., 2002). Indeed, the same authors performed VS simulation searching for antiviral drugs in an in-house library of compounds (Jin et al., 2020). Many other structures have been resolved since then, and there are several PDB entries with inhibitors located in the active site of the M^pro^ target (i.e. codes 5RG1, 5RGL, 7A1U, 7JU7) (Douangamath et al., 2020; Günther et al., 2021). Most of VS works strives for small molecules to reach the active site, which is characterized by a catalytic dyad of Cys145 and His41 residues (Zhang et al., 2020).

The SARS-CoV-2 spike has been also extensively assessed through VS simulations as it governs host attachment and virus–cell membrane fusion upon infection (Wu et al., 2020). In that framework, the characterization of the receptor-binding domain (RBD), deposited with PDB code 6M0J by Wang et al., 2020, allows for developing inhibitors by using molecular models (Tai et al., 2020). It has been also proposed that the Spike entity might evade antibody immunity by recruiting two metabolites, e.g., biliverdin and bilirubin (Rosa et al., 2021). Those allosteric sites are localized on the spike N-terminal domain (PDB code 7NTA).

All the crystallographic efforts devoted over the last two years have led to other X-ray structures, for instance for papain-like protease (PLpro), an enzyme that regulates SARS-CoV-2 viral spread and innate immunity (Shin et al., 2020). Some of the more recent crystals with non-covalent inhibitors are associated to the PDB codes 7CMD, 7JIR, 7JIT, 7JIV, 7JIW, 7JN2, 7JRN, 7KOJ, 7KOK, 7KOL, 7KRX, 7LBR, 7LBS, 7LLF, 7LLZ and 7LOS (Gao et al., 2021; Osipiuk et al., 2022, 2021; Ratia et al., 2014; Shen et al., 2021). In addition, the non-structural protein 16 (NSP16) has been shown to play an essential role for immune evasion by mimicking the human homolog, CMTr1. However, unlike CMTr1, NSP16 needs to form a heterodimer with NSP10 to activate its enzymatic activity. Although there are not available crystals for the inactive monomeric NSP16, Bowman and co-workers determined its activation mechanism and the location of a cryptic pocket (Vithani et al., 2021), that if targeted with a small molecule, can be used to inhibit NSP16.

It should be stressed a novel approach for reducing SARS-CoV-2 progression. Niemann-Pick type C1 (NPC1), a lipid-transfer protein that regulates intracellular cholesterol traffic, has been shown to play a role in human cell infection (García-Dorival et al., 2021). Within that hypothesis, the regulation of cholesterol by targeting NPC1 might offer an additional therapeutic strategy to treat infected patients. Unless viral targets, the experimental data for NPC1 is still scarce, with only two X-ray structures being available (PDB codes 6UOX and 5U73).

#### Library preparation, protein refinement and virtual screening

Our in-house focused library of 116 compounds were prepared for simulations by using LigPrep (Schrödinger Release 2021-3). Initial cartesian coordinates were retrieved from the PubChem data base (Kim et al., 2021). Geometries were subsequently optimized by assessing protonation and tautomeric states at pH 7±2 as predicted by Epik (Greenwood et al., 2010; Shelley et al., 2007). All other parameters are set as default as implemented in the Schrödinger suite of programs (Schrödinger Release 2021-3). All abovementioned PDB structures were refined with the Protein Preparation Wizard module (Madhavi Sastry et al., 2013), a multistep protocol. During that structural refinement, missing hydrogen atoms are added to the protein, charges were assigned, protonation states were determined with Epik at pH 7.0 ± 2.0, and a final restricted minimization is performed with the OPLS4 force field (Lu et al., 2021). A grid was created by considering co-crystal ligand or at the reference residues, as in the case of allosteric sites. Two docking runs were carried out, with single-precision (SP) or extra-precision (XP) scoring functions (Friesner et al., 2006; Halgren et al., 2004). For the records, ten poses per ligand were saved for each docking run. Binding energies are eventually computed with the molecular mechanics/generalized Born surface area (MMGBSA) implemented in Prime module (Jacobson et al., 2004, 2002), a more refined method that accounts for the energies before and after ligand binding (Rastelli et al., 2009; Ylilauri and Pentikäinen, 2013). The resulting free energies (ΔG) are therefore used for ranking compounds of our library. Last computational step uses molecular dynamics (MD) to assess the dynamic stability of the generated target-drug adducts. An orthorhombic box of TIP3P-waters was generated by using periodic boundary conditions with a buffer distance of 10 Å. Sodium cations are added to keep the system electronically neutral while additional ions were incorporated to mimic the physiological salt concentration of 0.15 M NaCl. The MD protocol includes minimization as implemented by default in the GPU-accelerated version of Desmond (Bowers et al., 2006). The MD simulation is completed by producing trajectories of 100 ns.

During our investigation a number of publications showed the antiviral activity of 10 compounds against SARS-CoV-2 (Simvastatin, Sirolimus, Everolimus, Itraconazole, Quercetin, Taxifolin, Resveratrol, Digoxin, Lanatoside C and SilibininA) that we found to inhibit 229E virus, and those were excluded from our study due to the lack of novelty (Cho et al., 2020; Jeon et al., 2020; Liesenborghs et al., 2021; Rodon et al., 2021; Van Damme et al., 2021). Indeed, some of them (Simvastatin, Sirolimus, Itraconazole, Quercetin, Resveratrol and Silibinin A) have been or are currently included in clinical trials for COVID-19 (https://clinicaltrials.gov).

### Reagents used

All the compounds and respective vendor origin are detailed in **Table 1**, and Suppl. Table 1.

**Table 1.**
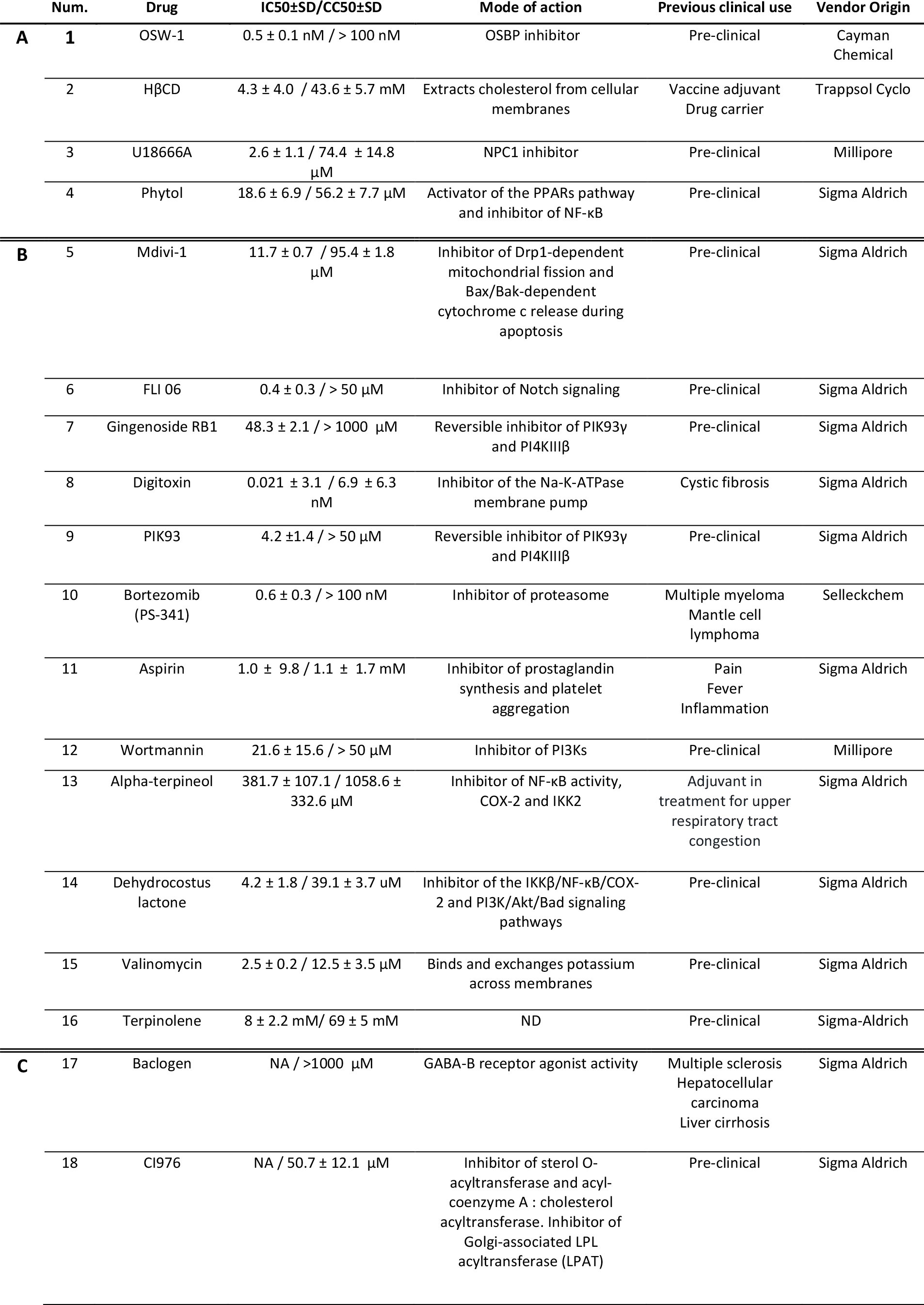
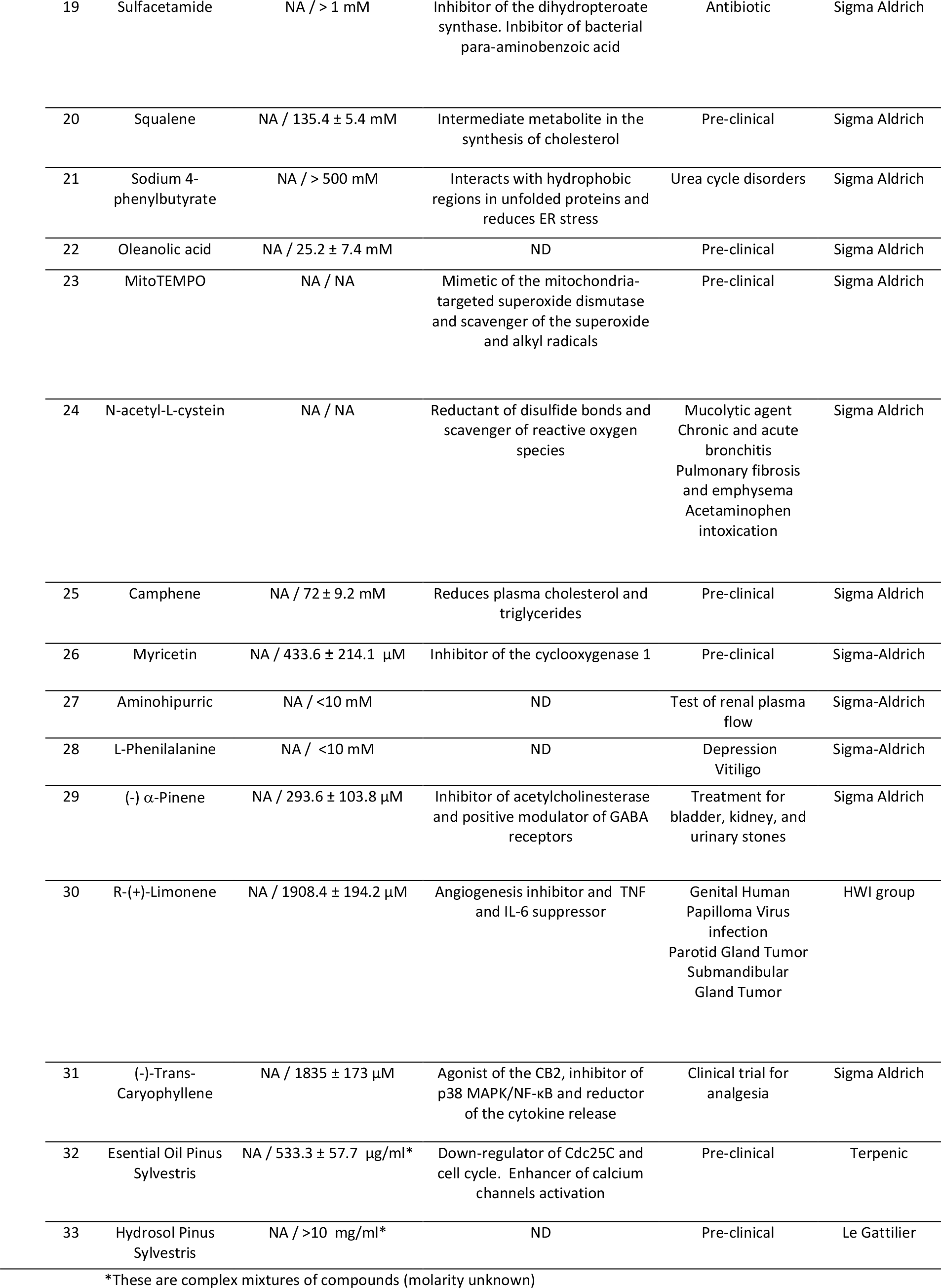
List of *in vitro* tested compounds against HCoV229E. The Table is divided in three sections: 1) Compounds with antiviral activity against HCoV229E and SARS-CoV-2 (A-white, 1-4); 2) compounds with antiviral activity against HCoV229E but not against SARS-CoV-2 (B-light grey, 5-16) and 3) compounds without antiviral activity against HCoV229E (C-dark grey, 17-33). For each compound, IC_50_ and CC_50_ values for HCoV229E, known mode of action, previous clinical use and vendor origin are indicated. ND, non-determined.

|  | Num. | Drug | IC50±SD/CC50±SD | Mode of action | Previous clinical use | Vendor Origin |
| --- | --- | --- | --- | --- | --- | --- |
| **A** | **1** | OSW-1 | 0.5 ± 0.1 nM / > 100 nM | OSBP inhibitor | Pre-clinical | Cayman Chemical |
|  | 2 | HβCD | 4.3 ± 4.0 / 43.6 ± 5.7 mM | Extracts cholesterol from cellular membranes | Vaccine adjuvant<br>Drug carrier | Trappsol Cyclo |
|  | 3 | U18666A | 2.6 ± 1.1 / 74.4 ± 14.8 μM | NPC1 inhibitor | Pre-clinical | Millipore |
|  | 4 | Phytol | 18.6 ± 6.9 / 56.2 ± 7.7 μM | Activator of the PPARs pathway and inhibitor of NF-κB | Pre-clinical | Sigma Aldrich |
| **B** | 5 | Mdivi-1 | 11.7 ± 0.7 / 95.4 ± 1.8 μM | Inhibitor of Drp1-dependent mitochondrial fission and Bax/Bak-dependent cytochrome c release during apoptosis | Pre-clinical | Sigma Aldrich |
|  | 6 | FLI 06 | 0.4 ± 0.3 / > 50 μM | Inhibitor of Notch signaling | Pre-clinical | Sigma Aldrich |
|  | 7 | Gingenoside RB1 | 48.3 ± 2.1 / > 1000 μM | Reversible inhibitor of PIK93γ and PI4KIIIβ | Pre-clinical | Sigma Aldrich |
|  | 8 | Digitoxin | 0.021 ± 3.1 / 6.9 ± 6.3 nM | Inhibitor of the Na-K-ATPase membrane pump | Cystic fibrosis | Sigma Aldrich |
|  | 9 | PIK93 | 4.2 ± 1.4 / > 50 μM | Reversible inhibitor of PIK93γ and PI4KIIIβ | Pre-clinical | Sigma Aldrich |
|  | 10 | Bortezomib (PS-341) | 0.6 ± 0.3 / > 100 nM | Inhibitor of proteasome | Multiple myeloma<br>Mantle cell lymphoma | Selleckchem |
|  | 11 | Aspirin | 1.0 ± 9.8 / 1.1 ± 1.7 mM | Inhibitor of prostaglandin synthesis and platelet aggregation | Pain<br>Fever<br>Inflammation | Sigma Aldrich |
|  | 12 | Wortmannin | 21.6 ± 15.6 / > 50 μM | Inhibitor of PI3Ks | Pre-clinical | Millipore |
|  | 13 | Alpha-terpineol | 381.7 ± 107.1 / 1058.6 ± 332.6 μM | Inhibitor of NF-κB activity, COX-2 and IKK2 | Adjuvant in treatment for upper respiratory tract congestion | Sigma Aldrich |
|  | 14 | Dehydrocostus lactone | 4.2 ± 1.8 / 39.1 ± 3.7 μM | Inhibitor of the IKKβ/NF-κB/COX-2 and PI3K/Akt/Bad signaling pathways | Pre-clinical | Sigma Aldrich |
|  | 15 | Valinomycin | 2.5 ± 0.2 / 12.5 ± 3.5 μM | Binds and exchanges potassium across membranes | Pre-clinical | Sigma Aldrich |
|  | 16 | Terpinolene | 8 ± 2.2 mM/ 69 ± 5 mM | ND | Pre-clinical | Sigma-Aldrich |
| **C** | 17 | Baclogen | NA / >1000 μM | GABA-B receptor agonist activity | Multiple sclerosis<br>Hepatocellular carcinoma<br>Liver cirrhosis | Sigma Aldrich |
|  | 18 | CI976 | NA / 50.7 ± 12.1 μM | Inhibitor of sterol O-acyltransferase and acyl-coenzyme A : cholesterol acyltransferase. Inhibitor of Golgi-associated LPL acyltransferase (LPAT) | Pre-clinical | Sigma Aldrich |

|  |  |  |  |  |  |
| --- | --- | --- | --- | --- | --- |
| 19 | Sulfacetamide | NA / > 1 mM | Inhibitor of the dihydropteroate synthase. Inhibitor of bacterial para-aminobenzoic acid | Antibiotic | Sigma Aldrich |
| 20 | Squalene | NA / 135.4 ± 5.4 mM | Intermediate metabolite in the synthesis of cholesterol | Pre-clinical | Sigma Aldrich |
| 21 | Sodium 4-phenylbutyrate | NA / > 500 mM | Interacts with hydrophobic regions in unfolded proteins and reduces ER stress | Urea cycle disorders | Sigma Aldrich |
| 22 | Oleanolic acid | NA / 25.2 ± 7.4 mM | ND | Pre-clinical | Sigma Aldrich |
| 23 | MitoTEMPO | NA / NA | Mimetic of the mitochondria-targeted superoxide dismutase and scavenger of the superoxide and alkyl radicals | Pre-clinical | Sigma Aldrich |
| 24 | N-acetyl-L-cystein | NA / NA | Reductant of disulfide bonds and scavenger of reactive oxygen species | Mucolytic agent<br>Chronic and acute bronchitis<br>Pulmonary fibrosis and emphysema<br>Acetaminophen intoxication | Sigma Aldrich |
| 25 | Camphene | NA / 72 ± 9.2 mM | Reduces plasma cholesterol and triglycerides | Pre-clinical | Sigma Aldrich |
| 26 | Myricetin | NA / 433.6 ± 214.1 µM | Inhibitor of the cyclooxygenase 1 | Pre-clinical | Sigma-Aldrich |
| 27 | Aminohipuric | NA / <10 mM | ND | Test of renal plasma flow | Sigma-Aldrich |
| 28 | L-Phenilalanine | NA / <10 mM | ND | Depression<br>Vitiligo | Sigma-Aldrich |
| 29 | (-) α-Pinene | NA / 293.6 ± 103.8 µM | Inhibitor of acetylcholinesterase and positive modulator of GABA receptors | Treatment for bladder, kidney, and urinary stones | Sigma Aldrich |
| 30 | R-(+)-Limonene | NA / 1908.4 ± 194.2 µM | Angiogenesis inhibitor and TNF and IL-6 suppressor | Genital Human Papilloma Virus infection<br>Parotid Gland Tumor<br>Submandibular Gland Tumor | HWI group |
| 31 | (-)-Trans-Caryophyllene | NA / 1835 ± 173 µM | Agonist of the CB2, inhibitor of p38 MAPK/NF-κB and reductor of the cytokine release | Clinical trial for analgesia | Sigma Aldrich |
| 32 | Esential Oil Pinus Sylvestris | NA / 533.3 ± 57.7 µg/ml* | Down-regulator of Cdc25C and cell cycle. Enhancer of calcium channels activation | Pre-clinical | Terpenic |
| 33 | Hydrosol Pinus Sylvestris | NA / >10 mg/ml* | ND | Pre-clinical | Le Gattilier |

\*These are complex mixtures of compounds (molarity unknown)

### Assays with HCoV-229E

#### Cells and virus

MRC5 cells (ATCC CCL-171) were grown in Dulbecco’s minimal essential medium (DMEM) supplemented with 10% inactivated fetal bovine serum (FBS, Biological Industries), 4 mM glutamine (Sigma-Aldrich), 1 x non-essential amino acid solution (Sigma-Aldrich), 100 U/ml penicillin and 100 μg/ml streptomycin (both from Sigma-Aldrich). Human coronavirus 229E (HCoV-229E; ATCC VR-740) was propagated in MRC-5 cells as described (Mesel-Lemoine et al., 2012) with a modification that is maintaining cell cultures at 35°C. Viral titer was calculated as 50% tissue culture infective dose (TCID50). Briefly, MRC-5 cells in a 96-well plate at 80% confluency were inoculated with a serial dilution of the viral stock, from 10^−1^ to 10^−8^. The plate was incubated at 33°C and with 5% CO_2_ during 5 days and then fixed with 4% paraformaldehyde (PFA) at room temperature (RT) during 20 minutes. Monolayers were processed by indirect immunofluorescence (IF) using specific antibodies against the HCoV-229E nucleocapsid (N) protein (see below), and titer calculated as described (Ramakrishnan, 2016; Reed and Muench, 1938).

#### Cytotoxicity Assay

A stock solution of each compound was prepared. Hydrophobic compounds were dissolved in dimethyl sulfoxide (DMSO) or ethanol (as recommended by the manufacturer) and stored at −20°C. To evaluate the viability of cell cultures when treated with the compounds, we performed an MTT (3-[4,5-dimethylthiazol-2-yl]-2,5 diphenyl tetrazolium bromide) assay, which measures the mitochondrial dehydrogenase activity of living cells (Shearman et al., 1994; Tolosa et al., 2015). MRC-5 cells were cultured in a 96-well plate until 80% confluency was reached. Serial dilutions of the compounds in DMEM with 10% FBS were added in triplicate and in a final volume of 100 μl/well. After 24 h, 5 mg/ml of MTT reagent (Sigma-Aldrich) was added to a final concentration of 0.5 mg/ml. Incubation was maintained during 4 h at 37°C with 5% CO_2_ for metabolization of the reagent and then 100 μl of lysis buffer (10 % SDS, 0.01 M HCl and 85% isopropanol) was added and incubation maintained for 30 minutes on an orbital shaker protected from light. Plates were read at 570 nm and the background (measured at 690 nm) was subtracted. Data ware calculated from three independent replicates.

#### Antiviral Activity

The effect of the compounds on viral infection was determined by IF as follows: 20,000 MRC5 cells/well were seeded in duplicates per drug and condition in 96-wells flat-bottom plates. When 80% confluency was reached, they were adsorbed with HCoV-229E in DMEM media without FBS at MOI= 0.1 PFU/cell, and incubated for 1h at 37°C. The inoculum was then removed and a serial dilution of the compounds diluted in DMEM supplemented with 1% FBS was added. A duplicate of the infected cells without any drug treatment served as control. After 24h, cells were fixed with 4% PFA for 20 min and washed three times with PBS. Cells were permeabilized with 0.25% saponin in PBS for 10 min and then treated 30 min with blocking buffer (1xPBS with 0.25% saponin and 2% FBS). Cells were then incubated 1 h with a rabbit antibody specific for the HCoV-229E nucleocapsid (N) protein (Ingenasa) diluted 1:200 in blocking buffer. After 3 washes with PBS, cells were incubated 45 min with an anti-rabbit secondary antibody conjugated with Alexa fluor 488 (Invitrogen) diluted 1:500 in blocking buffer and washed three times with PBS. Finally, cell nuclei were labeled 20 min with 4′,6-diamidino-2-phenylindole (DAPI) diluted 1:200 in blocking buffer and cells then washed 3 times with PBS. Images were obtained with a Leica DMi8 S widefield epifluorescence microscope and processed with Image J software. Data were normalized by setting the positive infection control as 100% of infection. Inhibition data were plotted as dose-effect curves fitted to a nonlinear regression model in GraphPad Prism v 9.4 software. The IC_50_ was calculated with Quest Graph™ IC50 Calculator (https://www.aatbio.com/tools/ic50-calculator). All experiments were replicated three times.

### Antiviral activity in cells infected with SARS-CoV-2

#### Biosafety statement

The biosafety committee of the Institute Germans Trias i Pujol approved the execution of SARS-CoV-2 experiments at the BSL3 laboratory of the Center for Bioimaging and Comparative Medicine (CSB-20-015-M7).

#### Cells

Vero E6 cells (ATCC CRL-1586) were cultured in Dulbecco’s modified Eagle medium, (DMEM) supplemented with 10% fetal bovine serum (FBS), 100 U/mL penicillin, 100 μg/mL streptomycin, and 2 mM glutamine (all from Invitrogen). HEK-293T (ATCC repository) were maintained in DMEM with 10% FBS, 100 IU/mL penicillin and 100 μg/mL streptomycin (all from Invitrogen). HEK-293T overexpressing the human ACE2 (293T-ACE2) were kindly provided by Integral Molecular Company and maintained in DMEM with 10% FBS, 100 IU/mL penicillin and 100 μg/mL streptomycin, and 1 μg/mL of puromycin (Invitrogen). CaLu-3 cells were kindly provided by the laboratory of Dr. Sanchez Cespedes and maintained in DMEM with 10% FBS, 100 IU/mL penicillin and 100 μg/mL streptomycin.

#### Virus isolation and sequencing

SARS-CoV-2 variants were isolated from clinical nasopharyngeal swabs in Vero E6 cells, as previously described (Rodon et al., 2021). Viral stocks were grown in Vero E6 cells and supernatants were collected and stored at −80°C until use. The following SARS-CoV-2 variants with deposited genomic sequence at the GISAID repository (http://gisaid.org) were tested: B.1 (D614G) isolated in Spain in March 2020 (EPI_ISL_510689); and 4 variants isolated in Spain from January to February 2021: Alpha or B.1.1.7 (EPI_ISL_1663569), ß or B.1.351 (originally detected in South Africa; EPI_ISL_1663571), Zeta or P.2 (originally detected in Brazil; EPI_ISL_1831696), Delta or B.1.617.2 (originally detected in India; EPI_ISL_3342900), and Omicron or B.1.1.529 (EPI_ISL_8151031). Genomic sequencing was performed from viral supernatant by using standard ARTIC v3 or v4 based protocols followed by Illumina sequencing [dx.doi.org/10.17504/protocols.io.bhjgj4jw]. Raw data analysis was performed by viralrecon pipeline [https://github.com/nf-core/viralrecon] while consensus sequence was called using samtools/ivar at the 75% frequency threshold. Viral variants were titrated at ½ dilutions on Vero E6 cells using the same luminometric assay described for antiviral testing. Thus, for all VOCs, we used equivalent infectious units inducing 50 % of viral induced cytopathic effect.

#### Pseudovirus production

HIV-1 reporter pseudovirus expressing SARS-CoV-2 Spike protein and luciferase were generated using two plasmids. pNL4-3.Luc.R-.E-was obtained from the NIH AIDS repository. SARS-CoV-2.SctΔ19 was generated (Geneart) from the full protein sequence of SARS-CoV-2 spike with a deletion of the last 19 amino acids in C-terminal, human-codon optimized and inserted into pcDNA3.4-TOPO (Ou et al., 2020). Spike plasmid was transfected with X-tremeGENE HP Transfection Reagent (Merck) into HEK-293T cells, and 24 hours post-transfection, cells were transfected with pNL4-3.Luc.R-.E-. Supernatants were harvested 48 hours later, filtered with 0.45 μM (Millex Millipore) and stored at −80°C until use. The p24gag content of all viruses was quantified using an ELISA (Perkin Elmer) and viruses were titrated in HEK-293T overexpressing the human ACE2.

#### Pseudoviral entry inhibition assay

HEK-293T overexpressing the human ACE2 were used to test the indicated compounds. A constant pseudoviral titer was used to pulse cells in the presence of the drugs. At 48 h post-inoculation, cells were lysed with the Bright Glo Luciferase system (Promega). Luminescence was measured with an EnSight Multimode Plate Reader (Perkin Elmer).

#### Antiviral activity against SARS-CoV-2

Increasing concentrations of the indicated antiviral compounds were added to Vero E6 cells and immediately after, we added equivalent infectious units of SARS-CoV-2 variants that induced a 50 % cytopathic effect. Untreated non-infected cells and untreated virus-infected cells were used as negative and positive controls of infection, respectively. To detect any drug-associated cytotoxic effect, Vero E6 cells were equally cultured in the presence of increasing drug concentrations, but in the absence of virus. Cytopathic or cytotoxic effects of the virus or drugs were measured 3 days after infection, using the CellTiter-Glo luminescent cell viability assay (Promega). Luminescence was measured in a Fluoroskan Ascent FL luminometer (ThermoFisher Scientific).

Viral replication of SARS-CoV-2 was also assessed on Calu-3 cells in the presence of the indicated antiviral compounds. Compounds were incubated with cells before adding the SARS-CoV-2 virus at MOI = 0.3. After 24 h of incubation at 37°C and 5% CO_2_, cells were washed with PBS and compounds were added in fresh media at the same concentration for 48h. The amount of SARS-CoV-2 nucleoprotein released to the supernatant was measured with SARS-CoV-2 nucleocapsid protein High-Sensitivity Quantitative ELISA (ImmunoDiagnostics) according to the manufacturer’s protocol. The cytopathic effect on Calu-3 cells was assessed with Cell Titer-Glo Assay with a Fluoroskan Ascent FL luminometer at the time of supernatant collection.

#### IC_50_ calculation and statistical analysis

Response curves of compounds or their mixes were adjusted to a non-linear fit regression model, calculated with a four-parameter logistic curve with variable slope. Cells not exposed to the virus were used as negative controls of infection, and were set as 100% of viability to normalize data and calculate the percentage of cytopathic effect. All analyses and Figures were generated with the GraphPad Prism v8.0b Software.

#### Transmission electron microscopy (TEM)

Mock- or SARS-CoV-2 – infected (MOI = 0.02 PFU/cell) cell monolayers were incubated in the absence or presence of two different concentrations of the compounds and fixed at 48 hpi with 4% paraformaldehyde and 1% glutaraldehyde in phosphate buffered saline (PBS) for 2h at room temperature (RT). Cells were removed from the plates in the fixative, pelleted by centrifugation and washed three times with PBS. Post-fixation of cell pellets was done on ice with 1% osmiumtetroxide + 0.8% potassium ferrocyanide in water. Afterwards, pellets were dehydrated on ice with increasing concentrations of acetone and processed for embedding in the epoxy resin EML-812 (Taab Laboratories), as previously described (Tenorio et al., 2018). Infiltration with epoxy resin was performed at RT. All samples were polymerized at 60°C for 48h. Ultrathin sections (50-70 nm) were cut with a Leica UC6 microtome and placed on uncoated 300 mesh copper grids. Sections were contrasted with 4% uranyl acetate and Reynold’s lead citrate. Images were taken with a Tecnai G2 TEM operated at 120kV with a Ceta camera or with a Jeol 1400 operated at 120kV with a Gatan Rio camera. At least 100 cells per condition were studied by TEM.

#### SARS-CoV-2 M^pro^ inhibition assay

A SARS-CoV-2 3CL main protease (MBP-tagged) assay kit (BPS Bioscience) was used following the kit instructions. In brief, the assay buffer was prepared by adding dithiothreitol, and then the protease was diluted to a 3–5 ng/μL. Next, diluted protease solution was added to the test samples and the positive controls. The MβCD and the GC376 control solutions were prepared with the assay buffer. The reaction was started by adding the substrate solution to each well and incubated overnight. Fluorescence was read in a EnSight multimode plate reader (Perkin Elmer) at a 360 nm/460 nm excitation/ detection wavelength. Percentage of M^pro^ activity inhibition is calculated as follows: (positive control – test inhibitor) / positive control.

#### Lipidomic analysis of plasma membranes of Calu-3 cells after MβCD treatment

To assess the lipid composition, 3 million of Calu-3 cells/well were seeded in a 6-well plate and treated with 0, 0.6 or 2.5 mM of MβCD for 2h at 37°C and 5% CO_2_. After extensive washing with PBS, cells were detached with trypsin and stored at −80°C until its analysis. A total of 750 μl of a methanol-chloroform (1:2, vol/vol) solution containing internal standards (16:0 D31_18:1 phosphocholine, C17:0 cholesteryl ester, stigmasterol and N-dodecanoylsphingosylphosphorylcholine, 0.2 nmol each, from Avanti Polar Lipids) were added to 3105cell. Samples were vortexed and sonicated until they appeared dispersed and extracted at 48°C overnight. The samples were then evaporated and transferred to 1.5 ml Eppendorf tubes after the addition of 0.5 ml of methanol. Samples were evaporated to dryness, and stored at −80°C until analysis. Before analysis, 150 μl of methanol were added to the samples, centrifuged at 13,000 g for 3 min, and 130 μl of the supernatants was transferred to ultra-performance liquid chromatography (UPLC) vials for injection and analysis with an Acquity UPLC system (Waters) connected to a time-of-flight (TOF; LCT Premier XE, Waters) detector (Simbari et al, 2016). Experiments were performed in duplicate and each sample was analyzed in triplicate.

## Results

### Molecular modeling for antiviral selection

We first screen the bibliography to identify: (i) molecules targeting RNA viral polymerases or cellular factors used by viruses; (ii) hydro-soluble extracts and essential oils from plants used in medicine and with antiviral activity; (iii) commercially available drugs that could have antiviral activity. These studies produced a library of 116 compounds that were all further analyzed with molecular models to identify potential inhibitors of SARS-CoV-2 proteins or cell proteins used by the virus. The designed computational strategy is summarized in **Figure 1**, including the selected viral targets (**Fig. 1A**) as well as the chemical structures of the best-ranked compounds (**Fig. 1B**). The performed calculations suggested that OSW-1 is efficiently anchored at the active site of the M^pro^ (**Fig. 1C**). Indeed, that hit exhibits the largest binding energy (ΔG = –108.79 kcal/mol) among all the small molecules included in our library. Additional large interactions are predicted for the binding of U18666A, wortmannin and phytol to NPC1 (ΔG in the range of –90 to –80 kcal/mol). Unfortunately, only moderate interactions are associated with other viral targets as Spike, NSP16 and PLPro, with ΔG values under the threshold of –66 kcal/mol. The complete list of the predicted binding-free energies for the best-ranked compounds is provided in the **Suppl. Table 2**.

**Figure 1.**
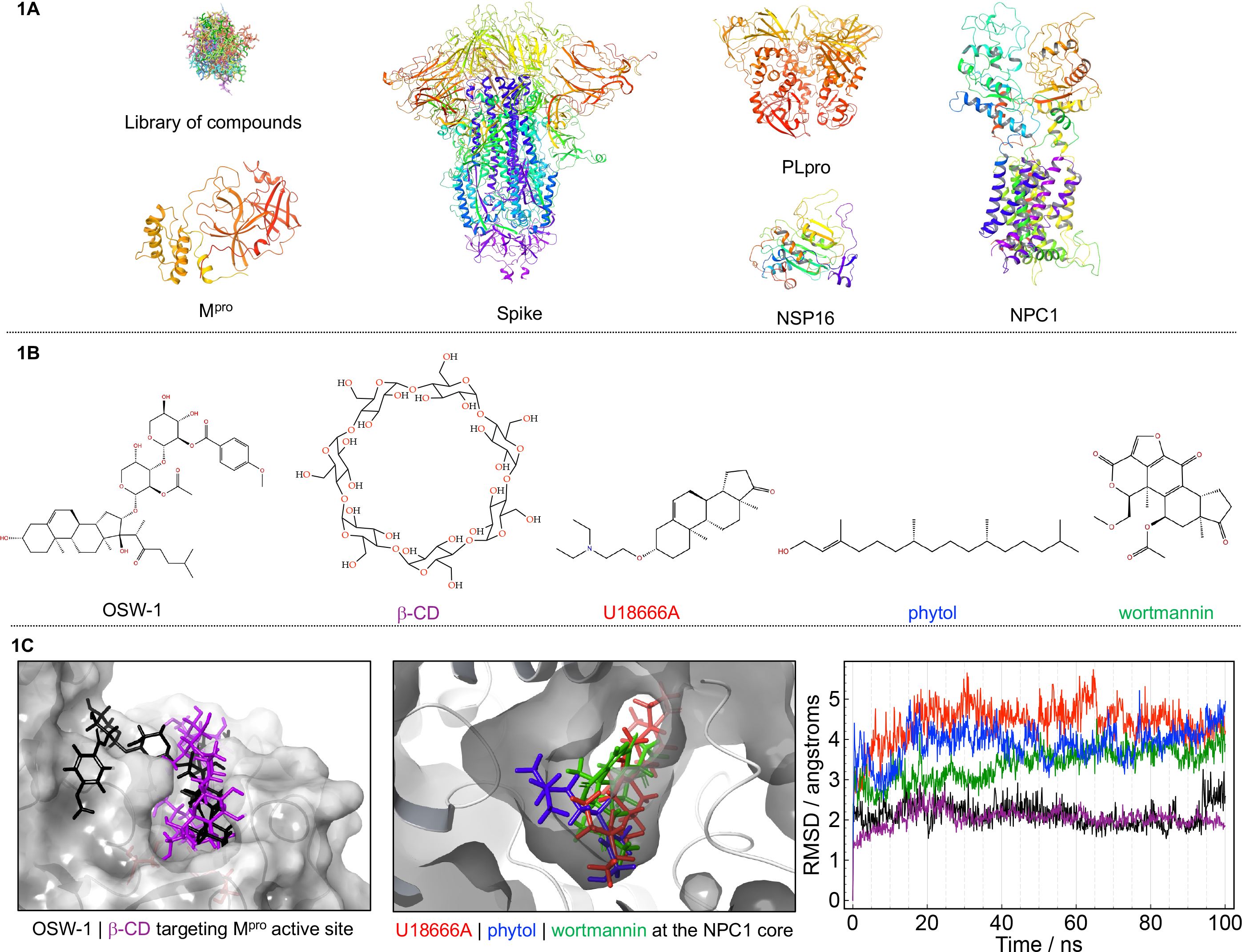
Virtual screening and molecular dynamics for selecting antiviral drugs. (**A**) Schematic representation of our library of compounds; experimental structures for all selected viral targets displayed in cartoons. (**B**) Chemical structures for the best-ranked compounds. (**C**) Poses adopted by the best-ranked compounds (targets are sketched in grey cartoons and surface); the associated root mean-square deviation (RMSD) in angstroms (Å). For the sake of clarity, the same color scheme has been adopted: black for OSW-1, red for U18666A, green for wortmannin, blue for phytol and purple for β-CD.

The adopted poses by the best-ranked ligands is illustrated in **Figure 1C** once reached the target. In the case of cyclodextrins, β-CD was selected as a representative macrocycle. The analysis of the produced structures confirms that both OSW-1 and β-CD are in the catalytic pocket of the M^pro^, while U1866A, phytol and wortmannin reach the central pocket of the NPC1. To ensure that ligands remain in their pockets under biological conditions, MD trajectories of 100 nano-seconds (ns) were performed. The stability was monitored through the so-called root means-square deviation (RMSD) (**Fig. 1C**, right panel). A close inspection of RMSD values revealed that both OSW-1 (black line) and β-CD (purple line) are stabilized with a RMSD variation of ca. 2Å. On the contrary, U18666A, phytol and wortmannin yielded an average RMSD of ca. 4Å. This difference is probably due to the fact that NCP1 has an internal pocket while the M^pro^ active site is localized in its surface, which induces a larger conformational change upon ligand binding. However, despite such dissimilarity, all five drugs were quickly equilibrated, with a variation of less than 1Å after the first 20ns of trajectory, suggesting the drug-target adducts are stable with time, a prerequisite to exhibit biological activity.

The energetic analysis offers an unbiased criterium for ligand selection. However, a large interaction with these targets does not guarantee an efficient activity against the virus, given that many other mechanisms may affect the inhibitory ability in the biological scenario (Cerón-Carrasco, 2022). Consequently, selection criteria for testing compounds in cell culture against coronaviruses HCoV-229E and SARS-CoV-2 was completed with additional parameters including their low toxicity and commercial availability. With this information, we selected 44 compounds to be tested first against HCoV-229E. In addition, we also tested the hydro-soluble extract and the essential oil from *Pinus sylvestris* in cells infected with HCoV-229E, given its described antiviral properties (Ha et al., 2020; Patel et al., 2016).

### Antiviral activity against HCoV-229E

To test the antiviral activity of selected compounds, MRC-5 cells were infected with HCoV-229E and treated in parallel with increasing concentrations of the selected compounds. Drug cytotoxicity was first measured, and safe drug-doses were used to determine the percentage of infected cells by immunofluorescence using an antibody specific for the HCoV-229E nucleocapsid protein (**Fig. 2A**). With these assays, we calculated the concentration of compound required to inhibit 50% of the virus (IC_50_), and the concentration for the 50% cytotoxic effect (CC_50_) (**Fig. 2B**). Thirteen compounds, including OSW-1, HP-β-cyclodextrin (HβCD), U18666A and Phytol inhibited HCoV-229E infection at non-toxic concentrations (**Table 1**, **Fig. 2B** top graphs). Mdivi-1, FLI06 and CI976 showed an inhibitory effect but at toxic concentrations, and Baclogen did not have an inhibitory effect on HCoV-229E (**Fig. 2B** bottom graphs). In the case of BMS309403, Dimercaprol, and alpha-cedrol, they inhibited 229E but their CC_50_ was close to the IC_50_ and they were discarded for further analysis due to toxicity issues. As indicated in **Table 1**, the most potent antiviral was OSW-1 (IC_50_ of 0.5nM), followed by U18666A, phytol and HβCD (IC_50_ of 2.6μM, 18.6 μM and 4.3mM, respectively).

**Figure 2.**
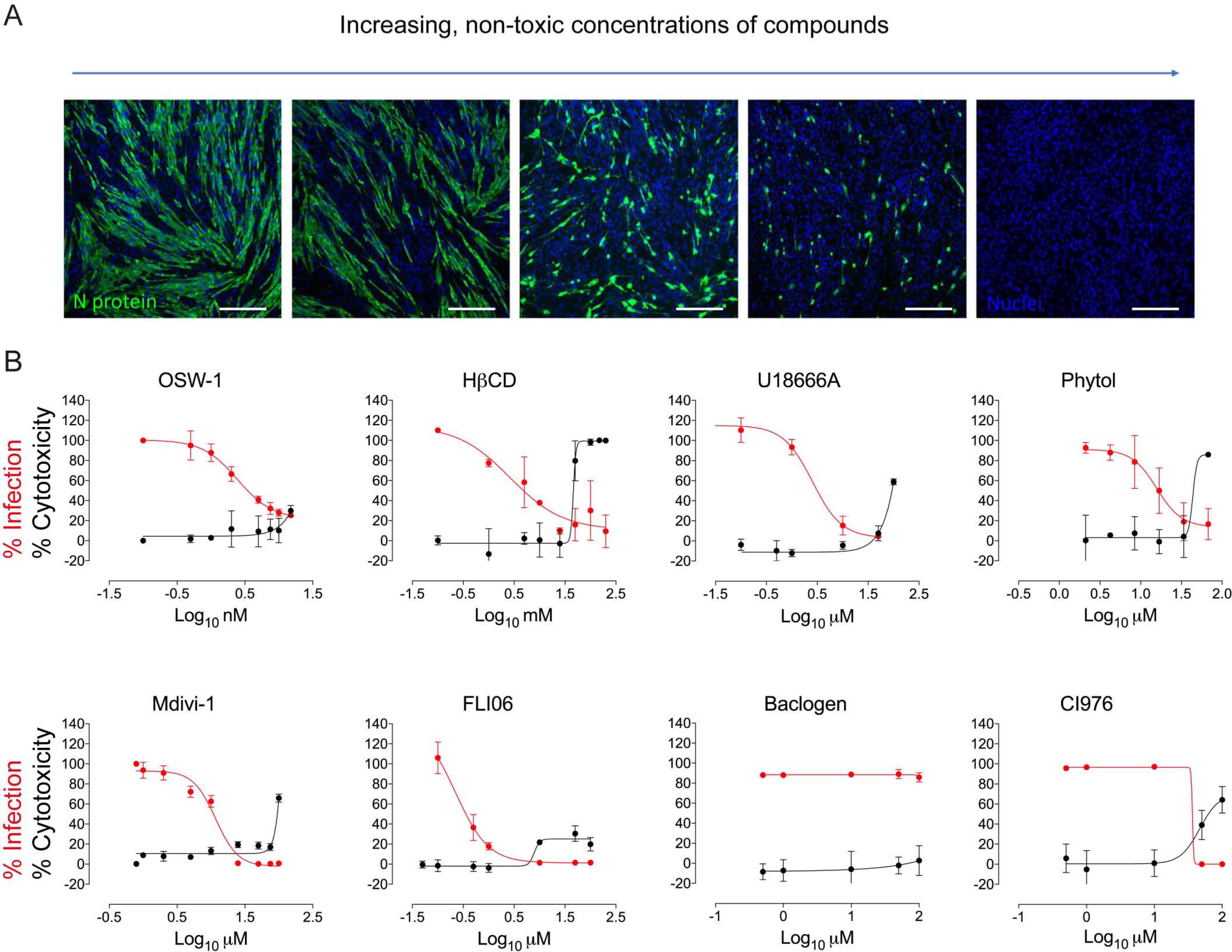
Antiviral activity of compounds tested against HCoV-229E. MRC-5 cells were absorbed with HCoV-229E at MOI of 0.1 PFUs/cell for 1h, exposed to increasing concentrations of the drug for 24h and processed by immunofluorescence with antibodies specific for the HCoV-229E N protein and with an anti-rabbit secondary antibody conjugated with Alexa fluor 488 (green). Nuclei were labeled with DAPI (blue). Images were collected with an epifluorescence microscope. (**A**) Representative pictures of the immunofluorescence assay. Scale bars, 100 μm. (**B**) Dose-response curves (red lines) of OSW-1, HβCD, U18666A, Phytol, Mdivi-1, FLI 06, Baclogen and CI976 were determined by nonlinear regression. Data is shown as mean ± S.E.M. of 3 biological replicates. Cytotoxic effect on MRC-5 cells exposed to increasing concentrations of drugs in the absence of virus is also shown (black lines).

### OSW-1, HβCD, U18666A and Phytol show anti-SARS-CoV-2 activity in Vero E6 cells

Compounds that revealed antiviral activity against HCoV-229E were next evaluated against SARS-CoV-2. Vero E6 cells were exposed to SARS-CoV-2 in the presence of increasing concentrations of the different drugs. After three days, the cytopathic effect of the virus and the cytotoxic effect of the drugs on cells were analyzed and the IC_50_ value calculated for each drug at non-cytotoxic concentrations. Results showed that OSW-1, HβCD, U18666A and Phytol had antiviral activity on these cells (**Fig. 3A**), while the rest of tested compounds were not active or were cytotoxic (**Fig. S1** and **Table 2**). The antiviral activity of these four drugs was similar when tested for different variants of concern of SARS-CoV-2 and comparable to the control Remdesivir (**Fig. 3B**). The antiviral activity found for tested compounds with both HCoV-229E and SARS-CoV-2 is summarized in **Table 1**.

**Table 2.**
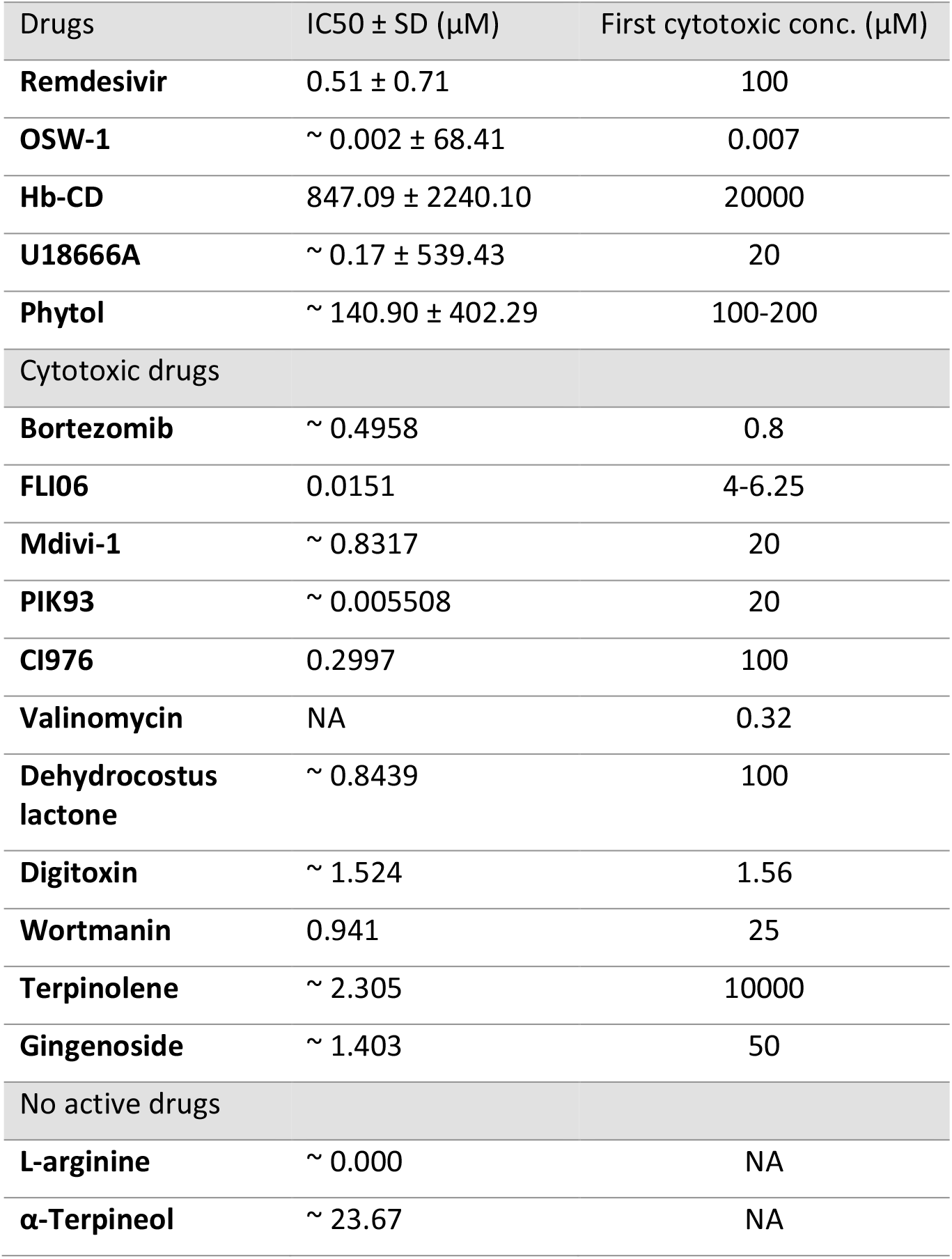
List of drugs tested in Vero E6 cells infected with SARS-CoV-2, for which the IC50 of the viral cytopathic effect is provided, or uninfected for which the first drug-cytotoxic concentration is provided. SD, standard deviation; NA, non-applicable.

**Figure 3.**
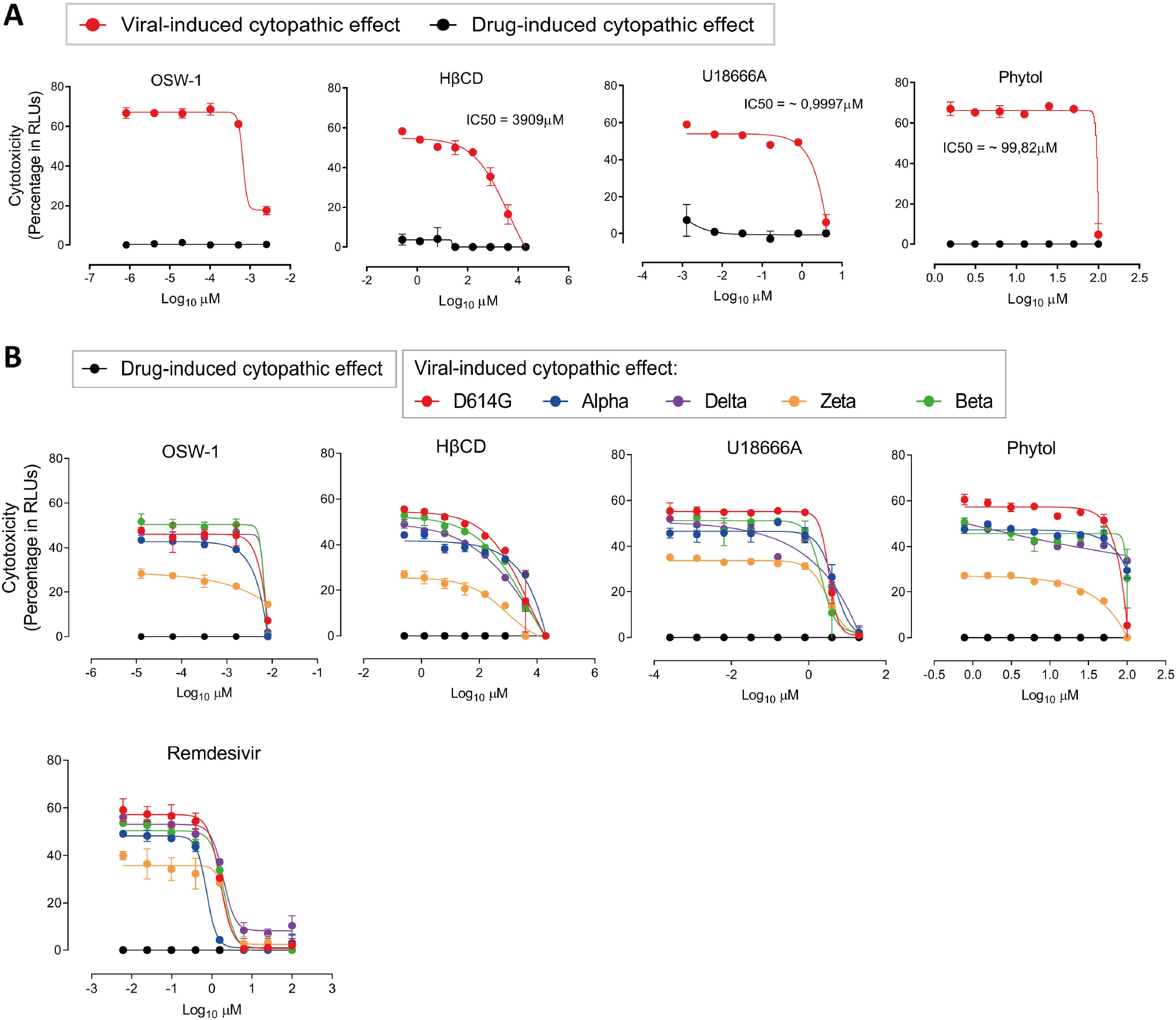
Antiviral activity of drugs against SARS-CoV-2. (**A**) Cytopathic effect on Vero E6 cells exposed to 200 TCID_50_/ml of SARS-CoV-2 in the presence of increasing concentrations of OSW-1 (1 – 1,3 x10^−5^ μM), HβCD (20 – 0,00026 mM), U18666A (20 – 2,6 x10^−4^ μM) and Phytol (100 – 0,78 μM). Non-linear fit to a variable response curve from one representative experiment out of three with two replicates is shown (red lines), excluding data from drug concentrations with associated toxicity. Cytotoxic effect with the same drug concentrations in the absence of virus is also shown (black lines). The IC_50_ value is indicated on each graph. IC_50_ for OSW-1 could not be calculated because 100 % inhibition was not obtained with this compound. (**B**) Cytopathic effect on Vero E6 cells exposed to 200 TCID_50_/ml of different variants of concern of SARS-CoV-2 as described in A, and in presence of Remdesivir.

Then we investigated the mechanism of action of the four selected antivirals active against α and β coronavirus. We first performed morphological studies of Vero E6 cells infected with SARS-CoV-2 by transmission electron microscopy (TEM). These studies showed characteristic structures assembled by SARS-CoV-2, which are: double-membrane vesicles (DMVs) where the virus replicates its genome (**Fig. 4A-C**), viral particles inside single-membrane vesicles (SMVs) (**Fig. 4D**), complex vacuoles (CV) with viruses (**Fig. 4E**) and extracellular viral particles (VPs) (**Fig. 4F**).

**Figure 4.**
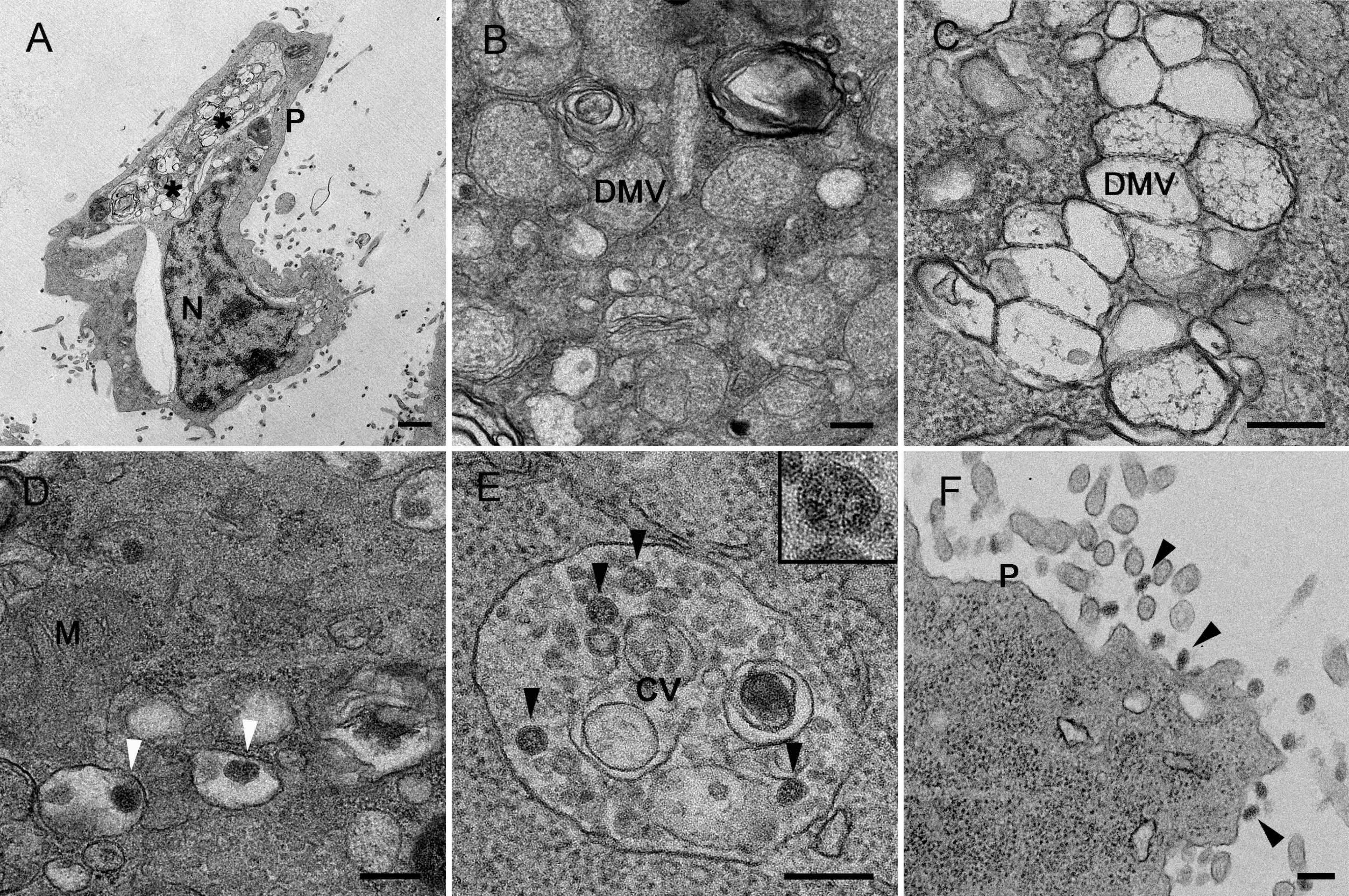
TEM of Vero E6 cells infected with SARS-CoV-2. Ultrathin sections of cells infected with SARS-CoV-2 at an MOI of 0.02 PFU/cell and 48 hpi. (**A**) Overview of a cell with clusters of DMVs (asterisks) near the nucleus (N). (**B**) High magnification of a group of early DMVs with electron-dense content. (**C**) Cluster of late DMVs with fibrillar content. (**D**) Single membrane vesicles with viral particles in their lumen (white arrowheads). (**E**) Complex vacuole (CV) with viral particles (black arrowheads) in their lumen. The inset shows a higher magnification of one of the viral particles inside the CV. (**F**) Viral particles (black arrowheads) at the plasma membrane (P). M, mitochondrion, Scale bars, 1 μm in A, 200 in B-F.

TEM of SARS-CoV-2 infected cells treated with two different concentrations of OSW-1 (**Fig. S2** and **Supp. Table 4**) and of U18666A (**Fig. S3** and **Supp. Table 4**) suggested that in Vero E6 cells, these compounds block DMVs’ assembly and function, which would affect subsequent steps of viral morphogenesis. Mock-infected cells incubated with OSW-1 exhibited normal morphology and no alterations were detected in any of the cell organelles compared to mock-infected cells in the absence of the drug (**Figure S4 and S5**). Incubation with the lower dose of U18666A caused no appreciable alterations in mock-infected cells (**Figure S6A-C**) but swelling of Golgi cisternae and lysosomes were observed when the higher dose is applied (**Figure S6D-F**). For phytol analysis, infected cells were incubated also with two different concentrations. Cells treated with the highest concentration showed clear signs of cytopathic effect and only the cells treated with the lowest concentration were processed for TEM. In phytol-treated cells, DMVs were altered to some extent but treatment had minor effects in virus assembly and egress (**Fig. S7A-E** and **Supp. Table 4**). In mock-infected cells, phytol did not cause significant alterations, with the only exception of mild increase in the amount of cytosolic glycogen granules and lipid droplets (**Supp. Fig. S7F-I**).

When SARS-CoV-2 exposed cells were treated with HβCD at the suboptimal concentration of 0.16 mM, DMVs showed alterations of the inner membrane (**Fig. 5A and B**). Viruses inside SMVs look normal, but large groups of distorted viral particles were seen inside complex vacuoles (**Fig 5C**). Extracellular virions attached to the plasma membrane looked normal (**Fig. 5D**). At inhibiting concentrations using 20 mM of HβCD, all viral structures were reduced (**Fig. 5E** and **Supp. Table 4**) and the few detected DMVs showed membrane alterations (**Fig. 5F and G**). Intracellular viral particles inside SMVs (**Fig. 5F**), CVs (**Fig. 5H**) and extracellular virions (**Fig. 5I**) were seen but in low numbers compared with infected non-treated cells (**Supp. Table 4**). These results show that HβCD affects the biogenesis of DMVs and the subsequent morphogenesis of new viral particles, an observation that could be linked to the capacity of this compound to reduce viral fusion and alter cellular membranes. At the lowest concentration, HβCD did not produce appreciable changes in cell compartments of mock-infected cells (**Figure S8A-C**). However, non-infected cells with 20mM of HβCD had increased number of vacuoles (**Fig. S8D**), swelling of ER, Golgi and lysosomes (**Fig. S8E and F**).

**Figure 5.**
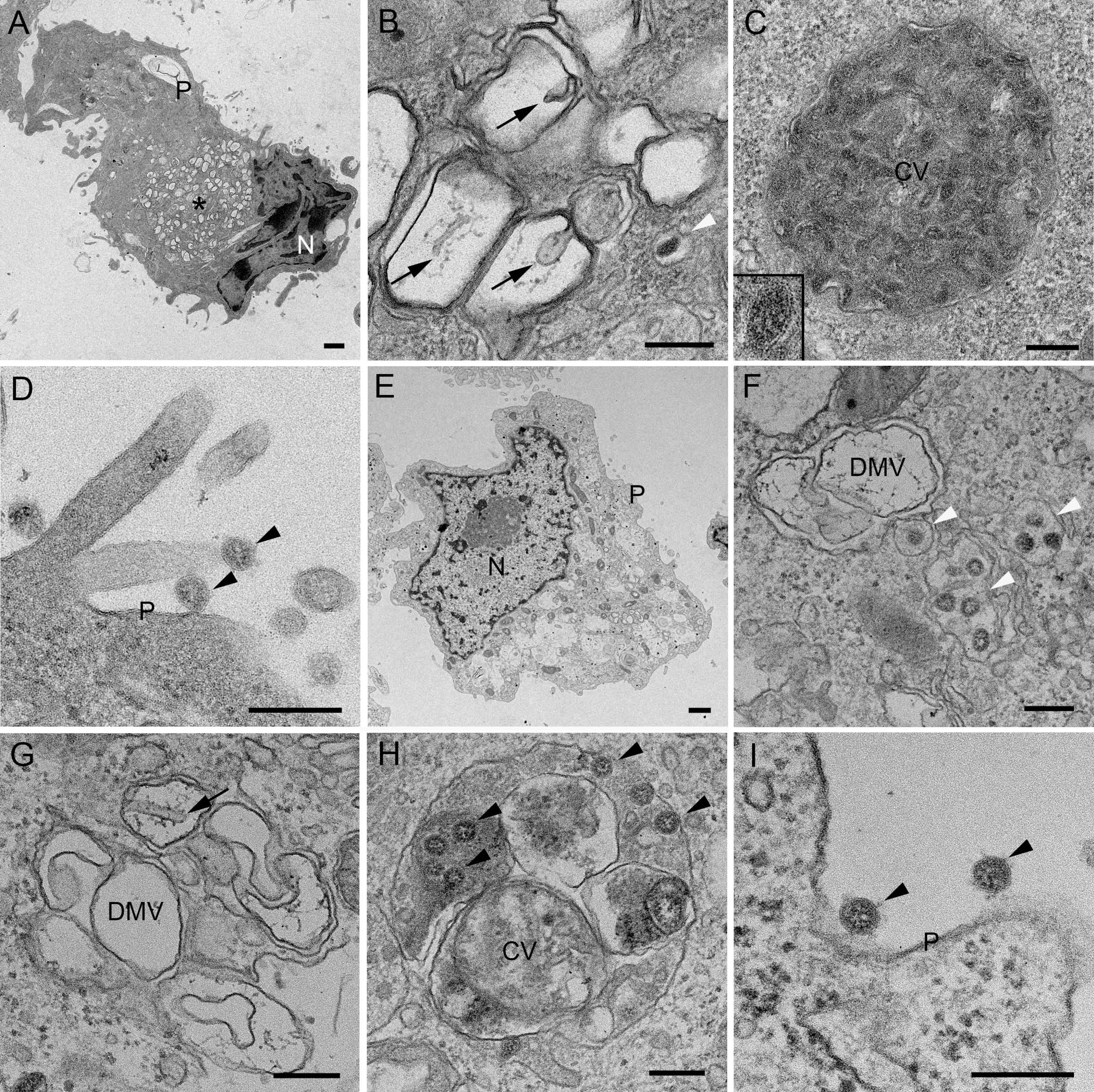
TEM of Vero E6 cells infected with SARS-CoV-2 and effects of HβCD. Cells infected with SARS-CoV-2 at an MOI of 0.02 PFU/cell were incubated with 0.16 mM (**A-D**) or 20 mM HβCD (**E-I**) and prepared for TEM at 48 hpi. (**A**) Overview of a cell with a cluster of DMVs (asterisk). (**B**) Group of DMVs with alteration of the inner membrane (arrows) and a single membrane vesicle with a viral particle (white arrowhead) in close vicinity. (**C**) Complex vacuole (CV) with viral particles in the lumen. The inset shows a viral particle at higher magnification. (**D**) Viral particles (black arrowheads) at the plasma membrane (P). (**E**) Overview of a cell. (**F**) DMV with a group of single membrane vesicles containing viral particles (white arrowheads) in close vicinity. (**G**) Group of DMVs with alteration of the inner membrane (arrow). (**H**) Complex vacuole with viral particles (arrowheads) in the lumen. (**I**) Viral particles (black arrowheads) at the plasma membrane (P). N, nucleus. Scale bars, 1 μm in A and E; 200 nm in B-D, F-I.

### HβCD and U18666A block viral fusion, but only HβCD inhibits infection in pulmonary cells

We therefore assessed the capability of these four drugs to inhibit the entry of SARS-CoV-2 pseudovirus in ACE2 expressing HEK-293T cells. Cells were exposed to fixed amounts of lentiviruses pseudotyped with SARS-CoV-2 spike in the presence of decreasing drug concentrations. Results showed that at drug concentrations without any cytotoxic effect, HβCD and U18666A inhibited the entry of pseudovirus, while OSW-1 and Phytol did not (**Fig. 6A)**. Cytopathic effect of the drugs in the absence of virus is shown **Fig. S9A**. These findings with the functional pseudoviral assay suggest the mechanism of action of cyclodextrins is reducing viral fusion, but it is also compatible with a possible interference with the proteolytic cleavage of M^Pro^ suggested by the molecular models.

**Figure 6.**
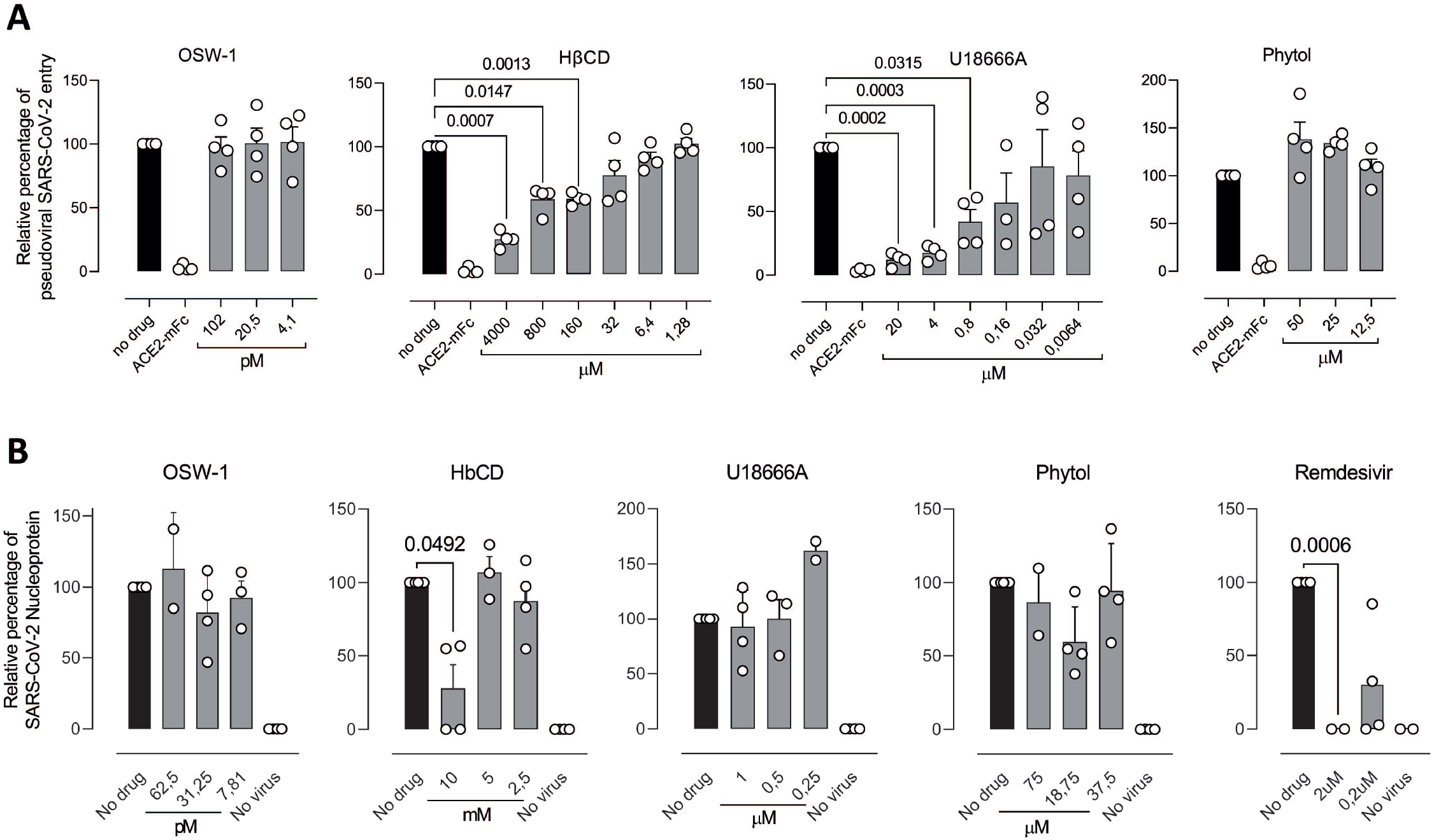
Drug inhibition of pseudovirus entry in ACE2-293T cells and of SARS-CoV-2 infection in pulmonary cells. (**A**) Relative viral entry of SARS-CoV-2 pseudoviruses in the presence of the indicated drugs in ACE2 expressing HEK-293T cells. Cells were exposed to fixed amounts of SARS-CoV-2 Spike lentiviruses in the presence of decreasing drug concentrations. Values show luciferase expression of the reporter lentiviruses pseudotyped with SARS-CoV-2, normalized to the luciferase expression of mock-treated cells (set at 100%). Mean and standard deviation from two experiments with two replicates each are represented, excluding cytotoxic values. (**B**) Relative viral replication of SARS-CoV-2 assessed on CaLu-3 cells in the presence of the indicated drugs. After 24h of adding virus and drugs at the indicated concentrations, cells were washed and compounds were added at the same final concentration for an additional 48h. Then supernatants were tested for viral release by detecting SARS-CoV-2 nucleocapsid concentration by ELISA. Values are normalized to the nucleocapsid concentration by mock-treated cells (set at 100%), which reached 5716 ± 2237 pg/mL (mean ± SD). Mean and standard deviation from four experiments are represented, excluding cytotoxic values.

We next tested SARS-CoV-2 antiviral activity in pulmonary Calu-3 cells as a more physiological and relevant cellular model. After incubating these cells with the drugs and the virus for 24h, the virus was washed away and compounds were added at the same concentration for 48h. The amount of SARS-CoV-2 nucleoprotein released to the supernatant was then measured by ELISA (**Fig. 6B**) and the cytopathic effect of the drugs in the absence of virus in Calu-3 cells was assessed by luminescence (**Fig. S9B**). Only HP-β-CD at 10 mM was able to effectively inhibit SARS-CoV-2 viral release into the supernatant of Calu-3 cells as observed for control remdesivir, while no inhibition was observed for the other compounds (**Fig. S9D**) at non-cytotoxic concentrations. Hence, HP-β-CD resulted as the most promising candidate to block SARS-CoV-2 replication on pulmonary cells.

### Other members of the Cyclodextrin family also inhibit SARS-CoV-2 infection in pulmonary cells

Given the well-known safety profiles of different types of β-cyclodextrins, that have been used as excipients in the pharmaceutical industry for decades (Lachowicz et al., 2020), we next aimed to test whether other cyclodextrins could hold the potential to inhibit viral replication. To test if other cyclodextrins could inhibit SARS-Cov-2 pseudoviral entry, we performed studies in ACE2 expressing HEK-293T cells. At no cytotoxic concentrations (**Fig. S9C**), seven cyclodextrins inhibited the pseudoviral SARS-CoV-2 entry, being the most potent β, HP-β, HP-γ and methyl-β-CDs (**Fig. 7A**).

**Figure 7.**
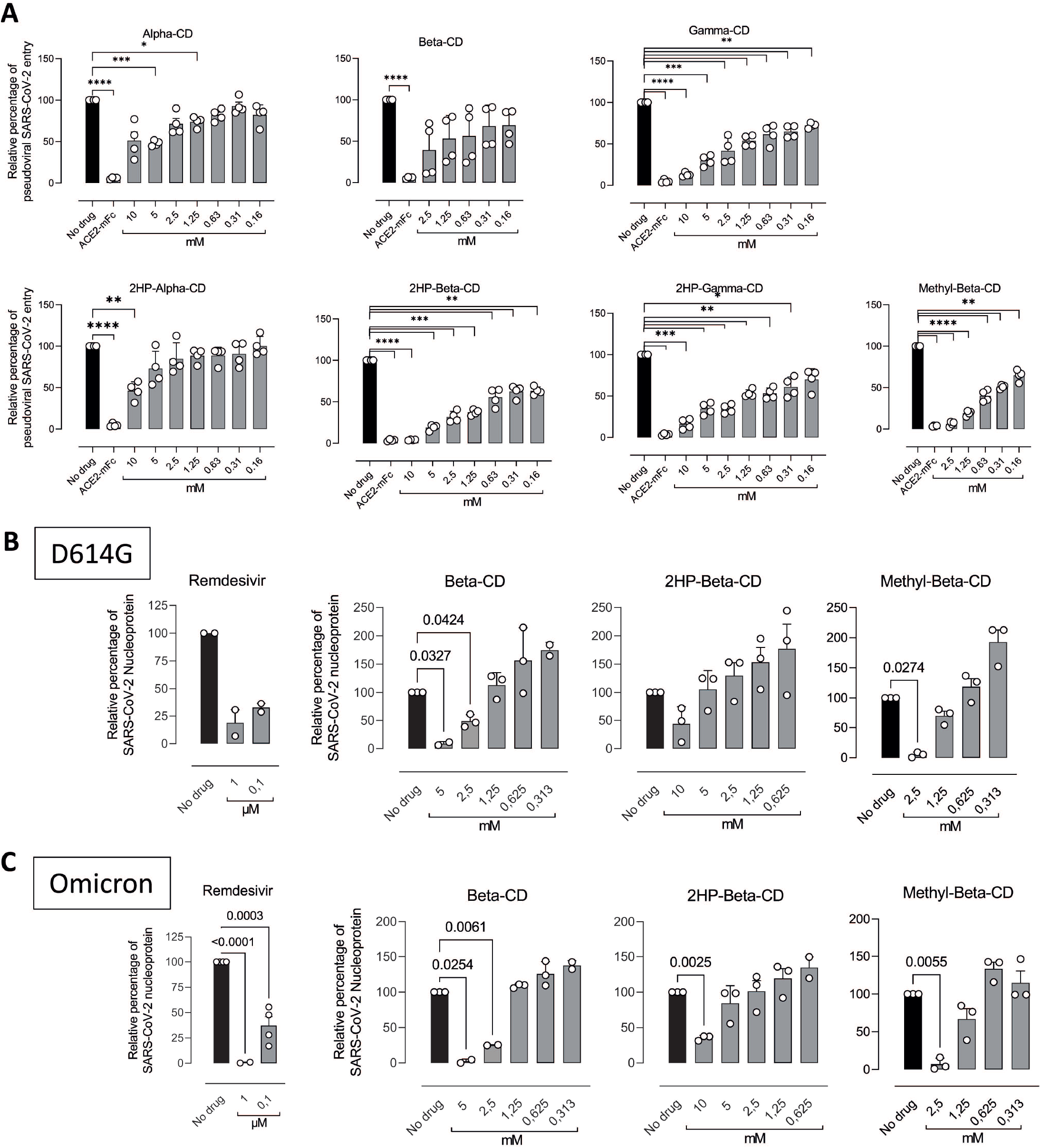
Members of the Cyclodextrin family inhibit SARS-CoV-2 infection in pulmonary cells. (**A**) Relative viral entry of SARS-CoV-2 pseudoviruses in the presence of the indicated cyclodextrins in ACE2 expressing HEK-293T cells. Cells were exposed to fixed amounts of SARS-CoV-2 Spike lentiviruses in the presence of decreasing drug concentrations. Values show luciferase expression of the reporter lentiviruses pseudotyped with SARS-CoV-2, normalized to the luciferase expression of mock-treated cells (set at 100%). Mean and standard deviation from two experiments with two replicates each are represented, excluding cytotoxic values. (**B**) Relative viral replication of D614G (**C**) or Omicron (**D**) SARS-CoV-2 variant was assessed on CaLu-3 cells in the presence of the indicated cyclodextrins. After 24h of adding virus and drugs at the indicated concentrations, cells were washed and compounds were added at the same final concentration for additional 48h. Then supernatants were tested for viral release by detecting SARS-CoV-2 nucleocapsid concentration by ELISA. Values are normalized to the nucleocapsid concentration by mock-treated cells (set at 100%), which reached 5716 ± 2237 pg/mL (mean ± SD). Mean and standard deviation from three experiments are represented, excluding cytotoxic values.

Next, we assayed if the most active cyclodextrins inhibiting pseudoviral fusion could also block SARS-CoV-2 replication in pulmonary Calu3 cells. The quantification of SARS-CoV-2 nucleoprotein released to the supernatant at no cytotoxic concentrations and detected by ELISA revealed that β-CDs, HP-β-CDs, and methyl-β-CDs were able to inhibit the SARS-CoV-2 viral activity on Calu-3 cells, both for the D614G (**Fig. 7B**) and for the Omicron BA.1 variant of concern (**Fig. 7C**). These results further highlight the potential of the β cyclodextrin family as antivirals against SARS-CoV-2.

### Cyclodextrins inhibit SARS-CoV-2 replication by interfering with viral fusion via cholesterol depletion

We had previously found that β-CD are in the catalytic pocket of the M^pro^ of SARS-CoV-2 by molecular modeling (Fig. 1C). This finding pointed to a possible explanation for the antiviral activity detected. All cyclodextrins, including α, β, γ, HP-α, HP-β, HP-γ and methyl-β-CDs, were screened by molecular modeling against the SARS-CoV-2 M^pro^. Since is possible to inhibit the action of M^pro^ by targeting two allosteric sites -PDB codes 7AXM and 7AGA-(Günther et al., 2021)), we expanded the chemical space of search into the designed models. Theory predicts that cyclodextrins can bind both active and allosteric M^pro^ sites, although significant dissimilarities appear depending on the size and nature of the macrocycle (**Suppl. Table 3**). To test whether this interaction occurred *in vitro*, the M^pro^ activity was measured in presence of increasing concentrations of methyl-β-CD (MβCD). While MβCD did not show activity, active GC376 control inhibited M^pro^ in a dose dependent manner (**Fig. 8A)**.

**Figure 8.**
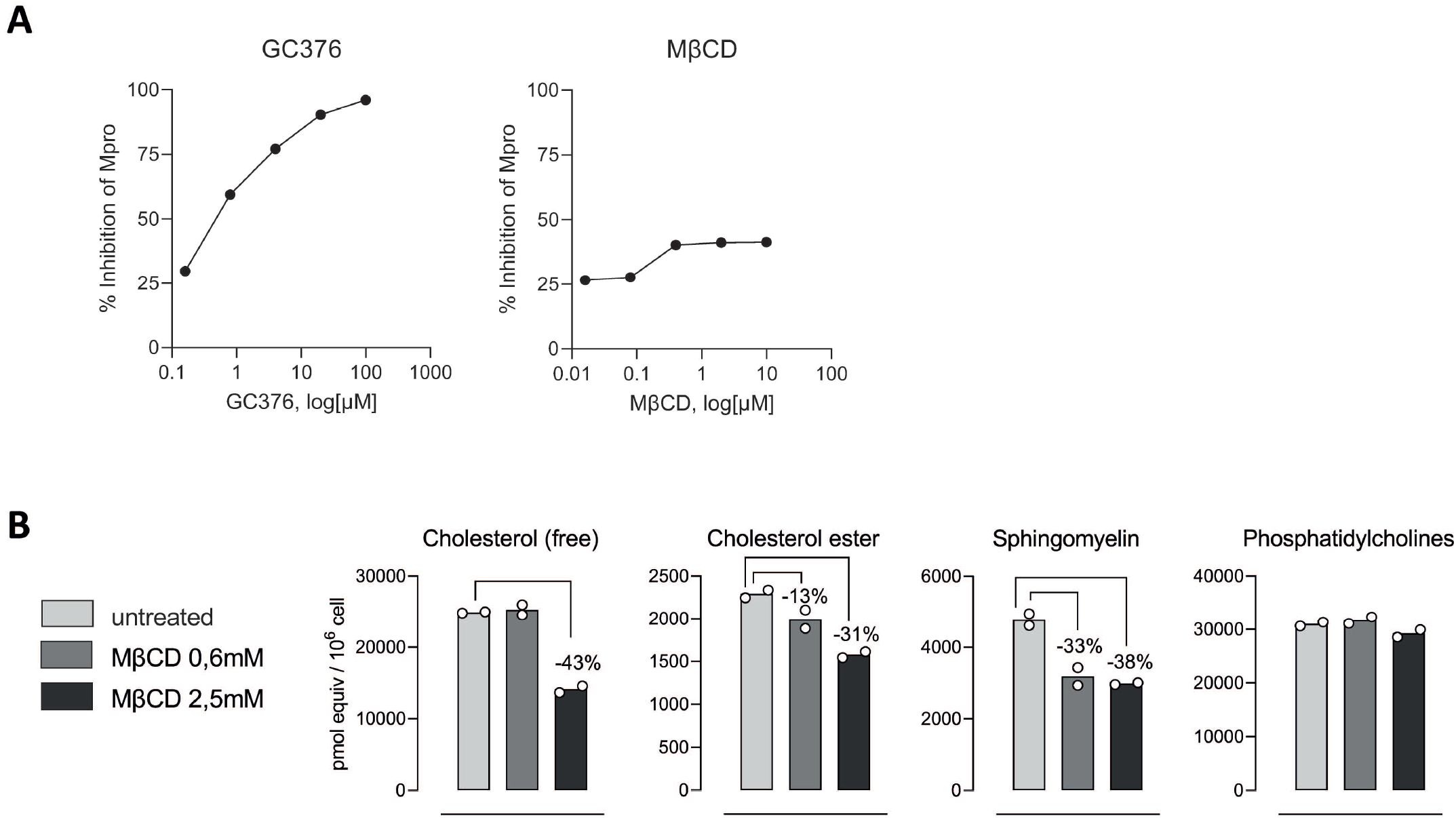
Cyclodextrins inhibit SARS-CoV-2 replication by interfering with viral fusion via cholesterol depletion. **(A)** M^pro^ activity measured in presence of increasing concentrations of methyl-β-CD (MβCD) or GC376 as positive inhibitor. Results are represented as the percentage of inhibition of M^pro^ activity in the absence of drugs. **(B)** Lipidomic measurement of plasma membranes from Calu3 cells treated or not with MβCD for 2h at 37°C and 5% CO_2_.

We next explored alternative antiviral mechanisms of action of cyclodextrins. Given the well-known capacity of cyclodextrins to extract cholesterol from biological membranes (López et al., 2011), we performed a lipidomic analysis focusing on different lipids associated to cholesterol enriched domains in biological membranes. Calu-3 cells treated with increasing concentrations of methyl-β-CDs showed a reduction of free and ester cholesterol along with sphingomyelin, and did not affect phosphatidylcholine (**Fig. 8B**). These results further confirm the capacity of cyclodextrins to alter the composition of cholesterol enriched domains actively involved in viral fusion processes. Taken together, these experiments along the pseudoviral fusion assays highlight the potential of cyclodextrins to inhibit SARS-CoV-2 replication by interfering with viral fusion via cholesterol depletion.

## DISCUSSION

Although pathogenic viruses pose a real and growing threat to public health, we have few medicines to prevent and treat viral infections. Here, a library of potential inhibitors of coronavirus infection was elaborated with a “based-on-knowledge” strategy. Examining the available information about what viruses use to complete their life cycle in cells and the description of the mechanism of action of drugs, we found compounds with potential to be used as antivirals to treat coronavirus infections. The main advantage of this strategy compared to high-throughput analysis is that the list of candidates is limited, and that different protocols of infection and drug treatment can be tested, which increases the probabilities of identifying molecules with antiviral activity. We have used a unique workflow involving biocomputational analysis and several biological assays to carefully select potential antivirals against different coronaviruses, including HCoV-229E and SARS-CoV-2. From our list of 116 compounds that target cell factors and pathways, 4 showed antiviral activity against both coronaviruses. Results with β-CDs were particularly relevant as showed consistent efficacy in different cellular models including human pulmonary cells. Particularly promising is our finding suggesting that the mechanism of action of β-CDs is interfering with viral fusion. Results from lipidomic analysis and transmission electron microscopy (TEM) showed that β-CDs may interfere with coronavirus infection by altering cholesterol and sphingomyelin content and disrupting the organization of membranes used by the virus. This is supported by our TEM results that showed dose-dependent effects of β-CDs in all SARS-CoV-2 structures in infected cells. The integrity of DMVs was compromised and viral morphogenesis was impaired, with the production of abnormal viral particles that are sometimes trapped in DMVs and inside large vacuoles that could represent degradation compartments. Our results confirm and broaden recent findings focused on HP-β-CDs (Bezerra et al., 2022), and with different methods further expand these results to other members of the cyclodextrin family and to SARS-CoV2 VOCs, including Omicron.

Given that cyclodextrins can be suited for oral, nasal or nebulized solutions, these results open different avenues to test diverse drug formulations that are known to be safe in humans (Stella and He, 2008; Tian et al., 2020). Further work should address the potential activity of these compounds in ameliorating SARS-CoV-2 infection in relevant animal models. The well-known safety profiles of β-CDs render these molecules as ideal candidates to develop affordable prophylactic and therapeutic compounds against coronaviruses (Fatmi et al., 2021; Sorice et al., 2020). Such drugs, which are already approved for clinical use in nasal spray devices (Guard et al., 1989; Paolacci et al., 2021), may be easier to deploy in low income countries compared to vaccines, which often require cold storage and must be administered by trained personnel. Given that β-CDs are widely used for compound encapsulation, they could be easily combined with other antivirals to potentiate activity and avoid viral resistance. Finally, the wide mechanism of action shown herein, which inhibits viral fusion with cellular membranes, could help to counteract other respiratory viruses, providing an arsenal to deploy in front of new variants of concern or future novel coronaviruses with pandemic potential. Broad-spectrum antivirals such as β-CDs could be ultimately applied to counteract unknown emergent viruses yet to appear, but need rigorous assessment in preclinical models for further development.

## ACKNOWLEDGEMENTS

Special thanks to Dr. Ma Carmen Martínez Jiménez, from the Hospital Nuestra Señora de Sonsoles de Avila for sharing knowledge about the use of plant extracts and their active compounds to treat viral infections.

## FUNDING

This work has been funded by grant RTI2018-094445-B100 (MCIU/AEI/FEDER, UE) from the Ministry of Science and Innovation of Spain (C.R.), by Palex Medical S.A., Sika S.A.U. and 7 more companies, and by Ms. Raquel Casaus Alvarez, Mr. Miguel Pardo Gil, Mr. Jacques Noguès and a total of 2,916 citizens through the Precipita crowdfunding platform of Fecyt (Fundación Española para la Ciencia y la Tecnología). NI-U is supported by the Spanish Ministry of Science and Innovation (grant PID2020-117145RB-I00), EU HORIZON-HLTH-2021-CORONA-01 (grant 101046118) and by institutional funding of Grifols, Pharma Mar, HIPRA, Amassence and Palobiofarma. This work used the computational resources of the Centro de Supercomputación de Galicia (CESGA) supported by the Partnership for Advanced Computing in Europe (PRACE) COVID-19 Fast Track Call for Proposals – Allocation Decision – Proposal COVID19-85.

## CONFLICT OF INTEREST

CR, IFC, MS, PO-G, AF-O, SF-S, NI-U, DP-Z, JM-B, DR-R, JPC-C, JAG and EN-D are inventors of a patent application for use of cyclodextrins to treat viral infection (US 63/323,091). The authors declare no other related competing interest.

## SUPPLEMENTARY MATERIALS

Table 2. List of *in vitro* **tested compounds against SARS-CoV-2**. The IC50 median and SDof the viral cytopathic effect from three independent experiments is provided, and the first drug-cytotoxic concentration is provided for uninfected cells. SD, standard deviation; NA, non-applicable.

**Suppl. Figure S1. Drugs without antiviral activity against SARS-CoV-2**. Cytopathic effect on Vero E6 cells exposed to 200 TCID_50_/ml of SARS-CoV-2 in the presence of increasing concentrations of different tested drugs. Non-linear fit to a variable response curve from one representative experiment out of three with two replicates is shown (red lines), excluding data from drug concentrations with associated toxicity. Cytotoxic effect with the same drug concentrations in the absence of virus is also shown (black lines).

**Suppl. Figure S2. TEM of Vero E6 cells infected with SARS-CoV-2 and effects of OSW-1**. Cells infected with SARS-CoV-2 at an MOI of 0.02 PFU/cell were incubated with 3 nM OSW-1 (**A-E**) or 5 nM OSW1 (**F-I**) and prepared for TEM at 48 hpi. (**A**) Overview of a cell with a cluster of DMVs (asterisk). (**B**) Detail of DMVs. (**C-D**) Examples for the variety of complex vacuoles (CV) with viral particles (arrowheads) in their lumen. The insets show higher magnifications of viral particles. (**E**) Viral particles (arrowheads) at the plasma membrane (P). (**F**) Overview of a cell with cluster of DMVs (asterisk). (**G**) Group of DMVs. The arrows mark alterations in their morphology such as protrusions in continuity with the inner membrane. The inset in (**G**) shows a viral particle between the inner and the outer membrane of the DMV. (**H**) Complex vacuole with viral particles in the lumen (arrowheads). The inset shows a viral particle at higher magnification. (**I**) Viral particles (arrowheads) at the plasma membrane (P). N, nucleus. Scale bars, 500 nm in A and F; 200 nm in B-E and G-I.

**Figure S3**. **TEM of Vero E6 cells infected with SARS-CoV-2 and effects of U18666A**. Cells were infected with SARS-CoV-2 at an MOI of 0.02 PFU/cell, incubated with 1 mM (**A-F**) or 20 mM U18666A (**G-I**) and prepared for TEM at 48 hpi. (**A**) Overview of a cell with a cluster of DMVs (asterisk). (**B**) Group of DMVs with alterations of the inner membrane (arrows). (**C**) Group of altered DMVs. In one of them, two viral particles (black arrowheads) are present between the inner and outer membrane of the DMV. Nearby there is a single membrane vesicle with a viral particle (white arrowhead). (**D**) Group of single membrane vesicles (white arrowheads) with viruses. (**E**) Complex vacuole with altered viral particles (arrowhead) in the lumen. The inset shows a viral particle at higher magnification. (**F**) Viral particles (arrowheads) at the plasma membrane (P). (**G**) Overview of a cell. (**H**) Micro-cluster of DMVs. (**I**) Viral particles (black arrowheads) at the plasma membrane. N, nucleus. Scale bars, 1 μm in A and G; 200 nm in B-F, H and I.

**Figure S4. TEM of mock-infected Vero E6 cells**. (**A**) Overview of a cell. (**B**) and (**C**) High magnification views of cells showing (**B**) a Golgi complex (G) and lysosomes (L) and (**C**) Mitochondria (M). (**D**) to (**F**) Details of (**D**) Nucleus (N), (**E**) Endoplasmatic reticulum (ER) and

(**F**) Plasma membrane (P). Scale bars, 1 μm in A, 200 nm in B-F.

**Figure S5. TEM of mock-infected Vero E6 cells treated with OSW-1**. Non-infected Vero cells were incubated with 3 nM (A-C) or 5 nM OSW1 (D-F) and prepared for TEM at 48 h post-incubation.(**A**) Overview of a cell. (**B**) and (**C**) High magnification views of cells showing a Golgi complex (G) and lysosomes (L). (**D**) Overview of a cell. (**E**) and (**F**) High magnification views showing a Golgi complex (G) and lysosomes (L). (**F**) Detail of a lysosome (L). N, nucleus. Scale bars, 500 nm in A and D; 200 nm in B, C, E and F.

**Figure S6. TEM of mock-infected Vero E6 cells treated with U18666A**. Cells were incubated with 1 mM (**A-C**) or 20 mM U18666A (**D-F**) and processed for TEM at 48 h post-incubation.(**A**) Overview of a cell. (**B**) and (**C**) High magnification views showing a Golgi complex (G) and a Lysosome (L). (**D**) Overview of a cell. (**E**) Swollen Golgi (G) with an endosomal vacuole (E) nearby. (**F**) Details of an enlarged lysosome (L). N, nucleus. Scale bars, 1 μm in A and D, 200 nm in B, C, E and F.

**Suppl. Figure S7. TEM of Vero E6 cells treated with Phytol and infected or not with SARS-CoV-2**. (**A-E**) Cells infected with SARS-CoV-2 at a MOI of 0.02 PFU/cell, incubated with 50 μM phytol and prepared for TEM at 48 hpi. (**A**) Overview of a cell with a cluster of DMVs (asterisk). (**B**) DMV with alteration of the inner membrane (arrow). (**C**) Group of single membrane vesicles with viral particles (white arrowheads). (**D**) Complex vacuole with viral particles in the lumen (arrowheads). The inset shows a viral particle at higher magnification. (**E**) Viral particles (black arrowheads) at the plasma membrane (P). N, nucleus. (**F-l**) Mock-infected cells were incubated with 50 μM phytol and prepared for TEM at 48 h post-incubation. (**F**) Overview of a cell. (**G**) Mitochondria (M). (**H**) Lysosome (L) and glycogen granules (white arrowheads) in the cytosol. (**I**) Lipid droplets (LD). P, plasma membrane. Scale bars, 1 μm in A and F, 200 nm in B-E, and G-I.

**Suppl. Figure S8. TEM of non-infected Vero E6 cells treated with HβCD**. Cells were incubated with 0.16 mM (**A-C**) or 20 mM HβCD (**D-F**) and prepared for TEM at 48 h post-incubation. (**A**) Overview of a cell. (**B**) and (**C**) High magnification views showing a Golgi complex (G) and Lysosomes (L). (**D**) Overview of a cell with vacuoles (V). (**E**) Detail of a Golgi complex with swollen cisternae (G) and swollen endoplasmatic reticulum (ER). (**F**) Details of enlarged lysosomes (L). N. nucleus. Scale bars, 1 μm in A and D, 200 nm in B, C, E and F.

**Figure S9. Cytotoxicity of drugs tested against SARS-CoV-2**. (**A**) Cytotoxicity of indicated drugs at decreasing concentrations in ACE2 expressing HEK-293T cells. At 48 h cells were lysed with the Glo Luciferase system (Promega) and luminescence was measured with a plate reader giving relative light units (RLUs). Cytotoxic concentrations are indicated with an arrow. Mean and standard deviation from one experiment with two replicates is represented. (**B**) Viral replication of SARS-CoV-2 assessed on Calu-3 cells in the presence of different drugs. After 24h of adding virus and drugs at the indicated concentrations, cells were washed and drugs were added at the same final concentration for an additional 48h. Then supernatants were tested for viral release by detecting SARS-CoV-2 nucleocapsid concentration by ELISA. (**C**) Cytotoxicity of indicated cyclodextrins at decreasing concentrations in ACE2 expressing HEK-293T cells. At 48 h cells were lysed with the Glo Luciferase system (Promega) and luminescence was measured with a plate reader giving relative light units (RLUs). Cytotoxic concentrations are indicated with an arrow. Mean and standard deviation from two experiments with two replicates each are represented. (**D**) Viral replication of SARS-CoV-2 assessed on Calu-3 cells in the presence of indicated cyclodextrins or remdesivir as control. After 24h of viral and drug addition, cells were washed and drugs were added at the same final concentration for an additional 48h. Then supernatants were tested for viral release by detecting SARS-CoV-2 nucleocapsid concentration by ELISA. Mean and standard deviation from three experiments are represented.

**Suppl. Table 1**. Vendor origin and concentrations of tested cyclodextrins.

**Suppl. Table 2**. Energies for the top-ranked compounds (small molecules) against Mpro, PLPro, NPC1, Spike, and NSP16.

**Suppl. Table 3**. Energies for the top-ranked cyclodextrins against the active and allosteric sites of Mpro.

**Suppl. Table 4**. Quantification of TEM results. Ultrathin sections of Vero E6 cells infected with SARS-CoV-2 at MOI of 0.02 PFU/cell and 48 hpi and treated in the absence or presence of OSW-1, HßCD, U18666A or Phytol were studied by TEM. The percentage of cells with DMVs, altered DMVs, single-membrane vesicles (SMVs) with viruses, complex vacuoles (CVs) with viruses and extracellular viral particles (VPs) was calculated. A total of 20 cells per condition were included in the quantification. Asterisk (*): very few DMVs per cell.

## Bibliography

Agarwal, A., Rochwerg, B., Lamontagne, F., Siemieniuk, R.A.C., Agoritsas, T., Askie, L., Lytvyn, L., Leo, Y.-S., Macdonald, H., Zeng, L., Amin, W., Barragan, F.A.J., Bausch, F.J., Burhan, E., Calfee, C.S., Cecconi, M., Chanda, D., Dat, V.Q., De Sutter, A., Du, B., Geduld, H., Gee, P., Harley, N., Hashmi, M., Hunt, B., Jehan, F., Kabra, S.K., Kanda, S., Kim, Y.-J., Kissoon, N., Krishna, S., Kuppalli, K., Kwizera, A., Lisboa, T., Mahaka, I., Manai, H., Mino, G., Nsutebu, E., Preller, J., Pshenichnaya, N., Qadir, N., Sabzwari, S., Sarin, R., Shankar-Hari, M., Sharland, M., Shen, Y., Ranganathan, S.S., Souza, J.P., Stegemann, M., Swanstrom, R., Ugarte, S., Venkatapuram, S., Vuyiseka, D., Wijewickrama, A., Maguire, B., Zeraatkar, D., Bartoszko, J.J., Ge, L., Brignardello-Petersen, R., Owen, A., Guyatt, G., Diaz, J., Kawano-Dourado, L., Jacobs, M., Vandvik, P.O., 2020. A living WHO guideline on drugs for covid-19. BMJ m3379. https://doi.org/10.1136/bmj.m3379

Benet, S., Blanch-Lombarte, O., Ainsua-Enrich, E., Pedreño-Lopez, N., Muñoz-Basagoiti, J., Raïch-Regué, D., Perez-Zsolt, D., Peña, R., Jiménez, E., de la Concepción, M.L.R., Ávila, C., Cedeño, S., Escribà, T., Romero-Martín, L., Alarcón-Soto, Y., Rodriguez-Lozano, G.F., Miranda, C., González, S., Bailón, L., Blanco, J., Massanella, M., Brander, C., Clotet, B., Paredes, R., Esteve, M., Izquierdo-Useros, N., Carrillo, J., Prado, J.G., Moltó, J., Mothe, B., 2022. Limited humoral and specific T-cell responses after SARS-CoV-2 vaccination in PLWH with poor immune reconstitution. J. Infect. Dis. https://doi.org/10.1093/infdis/jiac406

Berman, H.M., Battistuz, T., Bhat, T.N., Bluhm, W.F., Bourne, P.E., Burkhardt, K., Feng, Z., Gilliland, G.L., Iype, L., Jain, S., Fagan, P., Marvin, J., Padilla, D., Ravichandran, V., Schneider, B., Thanki, N., Weissig, H., Westbrook, J.D., Zardecki, C., 2002. The Protein Data Bank. Acta Crystallogr. Sect. D Biol. Crystallogr. 58, 899–907. https://doi.org/10.1107/S0907444902003451

Bezerra, B.B., Silva, G.P.D. da, Coelho, S.V.A., Correa, I.A., Souza, M.R.M. de, Macedo, K.V.G., Matos, B.M., Tanuri, A., Matassoli, F.L., Costa, L.J. da, Hildreth, J.E.K., Arruda, L.B. de, 2022. Hydroxypropyl-beta-cyclodextrin (HP-BCD) inhibits SARS-CoV-2 replication and virus-induced inflammatory cytokines. Antiviral Res. 205, 105373. https://doi.org/10.1016/j.antiviral.2022.105373

Bowers, K.J., Chow, E., Xu, H., Dror, R.O., Eastwood, M.P., Gregersen, B.A., Klepeis, J.L., Istvan Kolossvary, M., Moraes, A., Sacerdoti, F.D., Salmon, J.K., Shan, Y., Shaw, D.E., 2006. Scalable algorithms for molecular dynamics simulations on commodity clusters., in: In Proceedings of the 2006 ACM/IEEE Conference on Supercomputing (SC’06). Association for Computing Machinery. https://doi.org/10.1145/1188455.1188544

Buxeraud, J., Faure, S., Fougere, É., 2022. Le nirmatrelvir-ritonavir (Paxlovid®), un traitement contre la Covid-19. Actual. Pharm. 61, 10–12. https://doi.org/10.1016/j.actpha.2022.05.002

Cerón-Carrasco, J.P., 2022. When Virtual Screening Yields Inactive Drugs: Dealing with False Theoretical Friends. ChemMedChem 17. https://doi.org/10.1002/cmdc.202200278

Cho, J., Lee, Y.J., Kim, J.H., Kim, S. il, Kim, S.S., Choi, B.-S., Choi, J.-H., 2020. Antiviral activity of digoxin and ouabain against SARS-CoV-2 infection and its implication for COVID-19. Sci. Rep. 10, 16200. https://doi.org/10.1038/s41598-020-72879-7

Douangamath, A., Fearon, D., Gehrtz, P., Krojer, T., Lukacik, P., Owen, C.D., Resnick, E., Strain-Damerell, C., Aimon, A., Ábrányi-Balogh, P., Brandão-Neto, J., Carbery, A., Davison, G., Dias, A., Downes, T.D., Dunnett, L., Fairhead, M., Firth, J.D., Jones, S.P., Keeley, A., Keserü, G.M., Klein, H.F., Martin, M.P., Noble, M.E.M., O’Brien, P., Powell, A., Reddi, R.N., Skyner, R., Snee, M., Waring, M.J., Wild, C., London, N., von Delft, F., Walsh, M.A., 2020. Crystallographic and electrophilic fragment screening of the SARS-CoV-2 main protease. Nat. Commun. 11, 5047. https://doi.org/10.1038/s41467-020-18709-w

Fatmi, S., Taouzinet, L., Skiba, M., Iguer-Ouada, M., 2021. The Use of Cyclodextrin or its Complexes as a Potential Treatment Against the 2019 Novel Coronavirus: A Mini-Review. Curr. Drug Deliv. 18, 382–386. https://doi.org/10.2174/1567201817666200917124241

Friesner, R.A., Murphy, R.B., Repasky, M.P., Frye, L.L., Greenwood, J.R., Halgren, T.A., Sanschagrin, P.C., Mainz, D.T., 2006. Extra Precision Glide: Docking and Scoring Incorporating a Model of Hydrophobic Enclosure for Protein−Ligand Complexes. J. Med. Chem. 49, 6177–6196. https://doi.org/10.1021/jm051256o

Gao, X., Qin, B., Chen, P., Zhu, K., Hou, P., Wojdyla, J.A., Wang, M., Cui, S., 2021. Crystal structure of SARS-CoV-2 papain-like protease. Acta Pharm. Sin. B 11, 237–245. https://doi.org/10.1016/j.apsb.2020.08.014

García-Dorival, I., Cuesta-Geijo, M.Á., Barrado-Gil, L., Galindo, I., Garaigorta, U., Urquiza, J., Puerto, A. del, Campillo, N.E., Martínez, A., Gastaminza, P., Gil, C., Alonso, C., 2021. Identification of Niemann-Pick C1 protein as a potential novel SARS-CoV-2 intracellular target. Antiviral Res. 194, 105167. https://doi.org/10.1016/j.antiviral.2021.105167

Greenwood, J.R., Calkins, D., Sullivan, A.P., Shelley, J.C., 2010. Towards the comprehensive, rapid, and accurate prediction of the favorable tautomeric states of drug-like molecules in aqueous solution. J. Comput. Aided. Mol. Des. 24, 591–604. https://doi.org/10.1007/s10822-010-9349-1

Guard, S., Watling, K.J., Watson, S.P., 1989. Characterisation of [3H]-senktide binding to NK3 tachykinin receptors in guinea-pig ileum and cerebral cortex. Br. J. Pharmacol. 98 Suppl, 798P.

Günther, Sebastian, Reinke, P.Y.A., Fernández-García, Y., Lieske, J., Lane, T.J., Ginn, H.M., Koua, F.H.M., Ehrt, C., Ewert, W., Oberthuer, D., Yefanov, O., Meier, S., Lorenzen, K., Krichel, B., Kopicki, J.-D., Gelisio, L., Brehm, W., Dunkel, I., Seychell, B., Gieseler, H., Norton-Baker, B., Escudero-Pérez, B., Domaracky, M., Saouane, S., Tolstikova, A., White, T.A., Hänle, A., Groessler, M., Fleckenstein, H., Trost, F., Galchenkova, M., Gevorkov, Y., Li, C., Awel, S., Peck, A., Barthelmess, M., Schlünzen, F., Lourdu Xavier, P., Werner, N., Andaleeb, H., Ullah, N., Falke, S., Srinivasan, V., França, B.A., Schwinzer, M., Brognaro, H., Rogers, C., Melo, D., Zaitseva-Doyle, J.J., Knoska, J., Peña-Murillo, G.E., Mashhour, A.R., Hennicke, V., Fischer, P., Hakanpää, J., Meyer, J., Gribbon, P., Ellinger, B., Kuzikov, M., Wolf, M., Beccari, A.R., Bourenkov, G., von Stetten, D., Pompidor, G., Bento, I., Panneerselvam, S., Karpics, I., Schneider, T.R., Garcia-Alai, M.M., Niebling, S., Günther, C., Schmidt, C., Schubert, R., Han, H., Boger, J., Monteiro, D.C.F., Zhang, L., Sun, X., Pletzer-Zelgert, J., Wollenhaupt, J., Feiler, C.G., Weiss, M.S., Schulz, E.-C., Mehrabi, P., Karničar, K., Usenik, A., Loboda, J., Tidow, H., Chari, A., Hilgenfeld, R., Uetrecht, C., Cox, R., Zaliani, A., Beck, T., Rarey, M., Günther, Stephan, Turk, D., Hinrichs, W., Chapman, H.N., Pearson, A.R., Betzel, C., Meents, A., 2021. X-ray screening identifies active site and allosteric inhibitors of SARS-CoV-2 main protease. Science (80-.). 372, 642–646. https://doi.org/10.1126/science.abf7945

Ha, T.K.Q., Lee, B.W., Nguyen, N.H., Cho, H.M., Venkatesan, T., Doan, T.P., Kim, E., Oh, W.K., 2020. Antiviral Activities of Compounds Isolated from Pinus densiflora (Pine Tree) against the Influenza A Virus. Biomolecules 10, 711. https://doi.org/10.3390/biom10050711

Halgren, T.A., Murphy, R.B., Friesner, R.A., Beard, H.S., Frye, L.L., Pollard, W.T., Banks, J.L., 2004. Glide: A New Approach for Rapid, Accurate Docking and Scoring. 2. Enrichment Factors in Database Screening. J. Med. Chem. 47, 1750–1759. https://doi.org/10.1021/jm030644s

Jacobson, M.P., Friesner, R.A., Xiang, Z., Honig, B., 2002. On the Role of the Crystal Environment in Determining Protein Side-chain Conformations. J. Mol. Biol. 320, 597–608. https://doi.org/10.1016/S0022-2836(02)00470-9

Jacobson, M.P., Pincus, D.L., Rapp, C.S., Day, T.J.F., Honig, B., Shaw, D.E., Friesner, R.A., 2004. A hierarchical approach to all-atom protein loop prediction. Proteins Struct. Funct. Bioinforma. 55, 351–367. https://doi.org/10.1002/prot.10613

Jayk Bernal, A., Gomes da Silva, M.M., Musungaie, D.B., Kovalchuk, E., Gonzalez, A., Delos Reyes, V., Martín-Quirós, A., Caraco, Y., Williams-Diaz, A., Brown, M.L., Du, J., Pedley, A., Assaid, C., Strizki, J., Grobler, J.A., Shamsuddin, H.H., Tipping, R., Wan, H., Paschke, A., Butterton, J.R., Johnson, M.G., De Anda, C., 2022. Molnupiravir for Oral Treatment of Covid-19 in Nonhospitalized Patients. N. Engl. J. Med. 386, 509–520. https://doi.org/10.1056/NEJMoa2116044

Jeon, S., Ko, M., Lee, J., Choi, I., Byun, S.Y., Park, S., Shum, D., Kim, S., 2020. Identification of Antiviral Drug Candidates against SARS-CoV-2 from FDA-Approved Drugs. Antimicrob. Agents Chemother. 64. https://doi.org/10.1128/AAC.00819-20

Jin, Z., Du, X., Xu, Y., Deng, Y., Liu, M., Zhao, Y., Zhang, B., Li, X., Zhang, L., Peng, C., Duan, Y., Yu, J., Wang, L., Yang, K., Liu, F., Jiang, R., Yang, Xinglou, You, T., Liu, Xiaoce, Yang, Xiuna, Bai, F., Liu, H., Liu, Xiang, Guddat, L.W., Xu, W., Xiao, G., Qin, C., Shi, Z., Jiang, H., Rao, Z., Yang, H., 2020. Structure of Mpro from SARS-CoV-2 and discovery of its inhibitors. Nature 582, 289–293. https://doi.org/10.1038/s41586-020-2223-y

Lachowicz, M., Stańczak, A., Kołodziejczyk, M., 2020. Characteristic of Cyclodextrins: Their Role and Use in the Pharmaceutical Technology. Curr. Drug Targets 21, 1495–1510. https://doi.org/10.2174/1389450121666200615150039

Liesenborghs, L., Spriet, I., Jochmans, D., Belmans, A., Gyselinck, I., Teuwen, L.-A., ter Horst, S., Dreesen, E., Geukens, T., Engelen, M.M., Landeloos, E., Geldhof, V., Ceunen, H., Debaveye, B., Vandenberk, B., Van der Linden, L., Jacobs, S., Langendries, L., Boudewijns, R., Do, T.N.D., Chiu, W., Wang, X., Zhang, X., Weynand, B., Vanassche, T., Devos, T., Meyfroidt, G., Janssens, W., Vos, R., Vermeersch, P., Wauters, J., Verbeke, G., De Munter, P., Kaptein, S.J.F., Rocha-Pereira, J., Delang, L., Van Wijngaerden, E., Neyts, J., Verhamme, P., 2021. Itraconazole for COVID-19: preclinical studies and a proof-of-concept randomized clinical trial. EBioMedicine 66, 103288. https://doi.org/10.1016/j.ebiom.2021.103288

López, C.A., de Vries, A.H., Marrink, S.J., 2011. Molecular Mechanism of Cyclodextrin Mediated Cholesterol Extraction. PLoS Comput. Biol. 7, e1002020. https://doi.org/10.1371/journal.pcbi.1002020

Lu, C., Wu, C., Ghoreishi, D., Chen, W., Wang, L., Damm, W., Ross, G.A., Dahlgren, M.K., Russell, E., Von Bargen, C.D., Abel, R., Friesner, R.A., Harder, E.D., 2021. OPLS4: Improving Force Field Accuracy on Challenging Regimes of Chemical Space. J. Chem. Theory Comput. 17, 4291–4300. https://doi.org/10.1021/acs.jctc.1c00302

Madhavi Sastry, G., Adzhigirey, M., Day, T., Annabhimoju, R., Sherman, W., 2013. Protein and ligand preparation: parameters, protocols, and influence on virtual screening enrichments. J. Comput. Aided. Mol. Des. 27, 221–234. https://doi.org/10.1007/s10822-013-9644-8

Mesel-Lemoine, M., Millet, J., Vidalain, P.-O., Law, H., Vabret, A., Lorin, V., Escriou, N., Albert, M.L., Nal, B., Tangy, F., 2012. A human coronavirus responsible for the common cold massively kills dendritic cells but not monocytes. J Virol 86, 7577–87. https://doi.org/10.1128/JVI.00269-12

Molina, J.-M., Ghosn, J., Assoumou, L., Delaugerre, C., Algarte-Genin, M., Pialoux, G., Katlama, C., Slama, L., Liegeon, G., Beniguel, L., Ohayon, M., Mouhim, H., Goldwirt, L., Spire, B., Loze, B., Surgers, L., Pavie, J., Lourenco, J., Ben-Mechlia, M., Le Mestre, S., Rojas-Castro, D., Costagliola, D., 2022. Daily and on-demand HIV pre-exposure prophylaxis with emtricitabine and tenofovir disoproxil (ANRS PREVENIR): a prospective observational cohort study. Lancet HIV 9, e554–e562. https://doi.org/10.1016/S2352-3018(22)00133-3

Osipiuk, J., Azizi, S.-A., Dvorkin, S., Endres, M., Jedrzejczak, R., Jones, K.A., Kang, S., Kathayat, R.S., Kim, Y., Lisnyak, V.G., Maki, S.L., Nicolaescu, V., Taylor, C.A., Tesar, C., Zhang, Y.-A., Zhou, Z., Randall, G., Michalska, K., Snyder, S.A., Dickinson, B.C., Joachimiak, A., 2021. Structure of papain-like protease from SARS-CoV-2 and its complexes with non-covalent inhibitors. Nat. Commun. 12, 743. https://doi.org/10.1038/s41467-021-21060-3

Osipiuk, J., Wydorski, P.M., Lanham, B.T., Tesar, C., Endres, M., Engle, E., Jedrzejczak, R., Mullapudi, V., Michalska, K., Fidelis, K., Fushman, D., Joachimiak, A., Joachimiak, L.A., 2022. Dual domain recognition determines SARS-CoV-2 PLpro selectivity for human ISG15 and K48-linked di-ubiquitin. bioRxiv Prepr. Serv. Biol. https://doi.org/10.1101/2021.09.15.460543

Ou, X., Liu, Y., Lei, X., Li, P., Mi, D., Ren, L., Guo, L., Guo, R., Chen, T., Hu, J., Xiang, Z., Mu, Z., Chen, X., Chen, J., Hu, K., Jin, Q., Wang, J., Qian, Z., 2020. Characterization of spike glycoprotein of SARS-CoV-2 on virus entry and its immune cross-reactivity with SARS-CoV. Nat. Commun. 11, 1620. https://doi.org/10.1038/s41467-020-15562-9

Paolacci, S., Ergoren, M.C., De Forni, D., Manara, E., Poddesu, B., Cugia, G., Dhuli, K., Camilleri, G., Tuncel, G., Kaya Suer, H., Sultanoglu, N., Sayan, M., Dundar, M., Beccari, T., Ceccarini, M.R., Gunsel, I.S., Dautaj, A., Sanlidag, T., Connelly, S.T., Tartaglia, G.M., Bertelli, M., 2021. In vitro and clinical studies on the efficacy of α-cyclodextrin and hydroxytyrosol against SARS-CoV-2 infection. Eur. Rev. Med. Pharmacol. Sci. 25, 81–89. https://doi.org/10.26355/eurrev_202112_27337

Patel, N.K., Jaiswal, G., Bhutani, K.K., 2016. A review on biological sources, chemistry and pharmacological activities of pinostrobin. Nat. Prod. Res. 30, 2017–2027. https://doi.org/10.1080/14786419.2015.1107556

Ramakrishnan, M.A., 2016. Determination of 50% endpoint titer using a simple formula. World J. Virol. 5, 85–6. https://doi.org/10.5501/wjv.v5.i2.85

Rastelli, G., Rio, A. Del, Degliesposti, G., Sgobba, M., 2009. Fast and accurate predictions of binding free energies using MM-PBSA and MM-GBSA. J. Comput. Chem. NA-NA. https://doi.org/10.1002/jcc.21372

Ratia, K., Kilianski, A., Baez-Santos, Y.M., Baker, S.C., Mesecar, A., 2014. Structural Basis for the Ubiquitin-Linkage Specificity and deISGylating activity of SARS-CoV papain-like protease. PLoS Pathog. 10, e1004113. https://doi.org/10.1371/journal.ppat.1004113

Reed, L.J., Muench, H., 1938. A simple method of estimating fifty per cent endpoint. Am. J. Hyg. 27, 493–497.

Rodon, J., Muñoz-Basagoiti, J., Perez-Zsolt, D., Noguera-Julian, M., Paredes, R., Mateu, L., Quiñones, C., Perez, C., Erkizia, I., Blanco, I., Valencia, A., Guallar, V., Carrillo, J., Blanco, J., Segalés, J., Clotet, B., Vergara-Alert, J., Izquierdo-Useros, N., 2021. Identification of Plitidepsin as Potent Inhibitor of SARS-CoV-2-Induced Cytopathic Effect After a Drug Repurposing Screen. Front. Pharmacol. 12, 646676. https://doi.org/10.3389/fphar.2021.646676

Rosa, A., Pye, V.E., Graham, C., Muir, L., Seow, J., Ng, K.W., Cook, N.J., Rees-Spear, C., Parker, E., dos Santos, M.S., Rosadas, C., Susana, A., Rhys, H., Nans, A., Masino, L., Roustan, C., Christodoulou, E., Ulferts, R., Wrobel, A.G., Short, C.-E., Fertleman, M., Sanders, R.W., Heaney, J., Spyer, M., Kjær, S., Riddell, A., Malim, M.H., Beale, R., MacRae, J.I., Taylor, G.P., Nastouli, E., van Gils, M.J., Rosenthal, P.B., Pizzato, M., McClure, M.O., Tedder, R.S., Kassiotis, G., McCoy, L.E., Doores, K.J., Cherepanov, P., 2021. SARS-CoV-2 can recruit a heme metabolite to evade antibody immunity. Sci. Adv. 7. https://doi.org/10.1126/sciadv.abg7607

Schäfer, A., Martinez, D.R., Won, J.J., Meganck, R.M., Moreira, F.R., Brown, A.J., Gully, K.L., Zweigart, M.R., Conrad, W.S., May, S.R., Dong, S., Kalla, R., Chun, K., Du Pont, V., Babusis, D., Tang, J., Murakami, E., Subramanian, R., Barrett, K.T., Bleier, B.J., Bannister, R., Feng, J.Y., Bilello, J.P., Cihlar, T., Mackman, R.L., Montgomery, S.A., Baric, R.S., Sheahan, T.P., 2022. Therapeutic treatment with an oral prodrug of the remdesivir parental nucleoside is protective against SARS-CoV-2 pathogenesis in mice. Sci. Transl. Med. 14. https://doi.org/10.1126/scitranslmed.abm3410

Shearman, M.S., Ragan, C.I., Iversen, L.L., 1994. Inhibition of PC12 cell redox activity is a specific, early indicator of the mechanism of beta-amyloid-mediated cell death. Proc. Natl. Acad. Sci. U. S. A. 91, 1470–4. https://doi.org/10.1073/pnas.91.4.1470

Shelley, J.C., Cholleti, A., Frye, L.L., Greenwood, J.R., Timlin, M.R., Uchimaya, M., 2007. Epik: a software program for pK a prediction and protonation state generation for drug-like molecules. J. Comput. Aided. Mol. Des. 21, 681–691. https://doi.org/10.1007/s10822-007-9133-z

Shen, Z., Ratia, K., Cooper, L., Kong, D., Lee, H., Kwon, Y., Li, Y., Alqarni, S., Huang, F., Dubrovskyi, O., Rong, L., Thatcher, G.R., Xiong, R., 2021. Potent, Novel SARS-CoV-2 PLpro Inhibitors Block Viral Replication in Monkey and Human Cell Cultures. bioRxiv Prepr. Serv. Biol. https://doi.org/10.1101/2021.02.13.431008

Shin, D., Mukherjee, R., Grewe, D., Bojkova, D., Baek, K., Bhattacharya, A., Schulz, L., Widera, M., Mehdipour, A.R., Tascher, G., Geurink, P.P., Wilhelm, A., van der Heden van Noort, G.J., Ovaa, H., Müller, S., Knobeloch, K.-P., Rajalingam, K., Schulman, B.A., Cinatl, J., Hummer, G., Ciesek, S., Dikic, I., 2020. Papain-like protease regulates SARS-CoV-2 viral spread and innate immunity. Nature 587, 657–662. https://doi.org/10.1038/s41586-020-2601-5

Sorice, M., Misasi, R., Riitano, G., Manganelli, V., Martellucci, S., Longo, A., Garofalo, T., Mattei, V., 2020. Targeting Lipid Rafts as a Strategy Against Coronavirus. Front. cell Dev. Biol. 8, 618296. https://doi.org/10.3389/fcell.2020.618296

Stella, V.J., He, Q., 2008. Cyclodextrins. Toxicol. Pathol. 36, 30–42. https://doi.org/10.1177/0192623307310945

Tai, W., He, L., Zhang, X., Pu, J., Voronin, D., Jiang, S., Zhou, Y., Du, L., 2020. Characterization of the receptor-binding domain (RBD) of 2019 novel coronavirus: implication for development of RBD protein as a viral attachment inhibitor and vaccine. Cell. Mol. Immunol. 17, 613–620. https://doi.org/10.1038/s41423-020-0400-4

Tenorio, R., de Castro, I.F., Knowlton, J.J., Zamora, P.F., Lee, C.H., Mainou, B.A., Dermody, T.S., Risco, C., 2018. Reovirus σNS and μNS Proteins Remodel the Endoplasmic Reticulum to Build Replication Neo-Organelles. MBio 9, 15.

Tian, B., Hua, S., Liu, J., 2020. Cyclodextrin-based delivery systems for chemotherapeutic anticancer drugs: A review. Carbohydr. Polym. 232, 115805. https://doi.org/10.1016/j.carbpol.2019.115805

Tolosa, L., Donato, M.T., Gómez-Lechón, M.J., 2015. General Cytotoxicity Assessment by Means of the MTT Assay. pp. 333–348. https://doi.org/10.1007/978-1-4939-2074-7_26

Van Damme, E., De Meyer, S., Bojkova, D., Ciesek, S., Cinatl, J., De Jonghe, S., Jochmans, D., Leyssen, P., Buyck, C., Neyts, J., Van Loock, M., 2021. In vitro activity of itraconazole against SARS-CoV-2. J. Med. Virol. 93, 4454–4460. https://doi.org/10.1002/jmv.26917

Vithani, N., Ward, M.D., Zimmerman, M.I., Novak, B., Borowsky, J.H., Singh, S., Bowman, G.R., 2021. SARS-CoV-2 Nsp16 activation mechanism and a cryptic pocket with pan-coronavirus antiviral potential. Biophys. J. 120, 2880–2889. https://doi.org/10.1016/j.bpj.2021.03.024

Wu, C., Liu, Y., Yang, Y., Zhang, P., Zhong, W., Wang, Y., Wang, Q., Xu, Y., Li, M., Li, X., Zheng, M., Chen, L., Li, H., 2020. Analysis of therapeutic targets for SARS-CoV-2 and discovery of potential drugs by computational methods. Acta Pharm. Sin. B 10, 766–788. https://doi.org/10.1016/j.apsb.2020.02.008

Yang, K.S., Leeuwon, S.Z., Xu, S., Liu, W.R., 2022. Evolutionary and Structural Insights about Potential SARS-CoV-2 Evasion of Nirmatrelvir. J. Med. Chem. 65, 8686–8698. https://doi.org/10.1021/acs.jmedchem.2c00404

Ylilauri, M., Pentikäinen, O.T., 2013. MMGBSA As a Tool To Understand the Binding Affinities of Filamin–Peptide Interactions. J. Chem. Inf. Model. 53, 2626–2633. https://doi.org/10.1021/ci4002475

Zhang, L., Lin, D., Sun, X., Curth, U., Drosten, C., Sauerhering, L., Becker, S., Rox, K., Hilgenfeld, R., 2020. Crystal structure of SARS-CoV-2 main protease provides a basis for design of improved α-ketoamide inhibitors. Science (80-.). 368, 409–412. https://doi.org/10.1126/science.abb3405

