## Supplementary_Material for "β-Cyclodextrins as affordable antivirals to treat coronavirus infection"

### Supplementary Figure 1

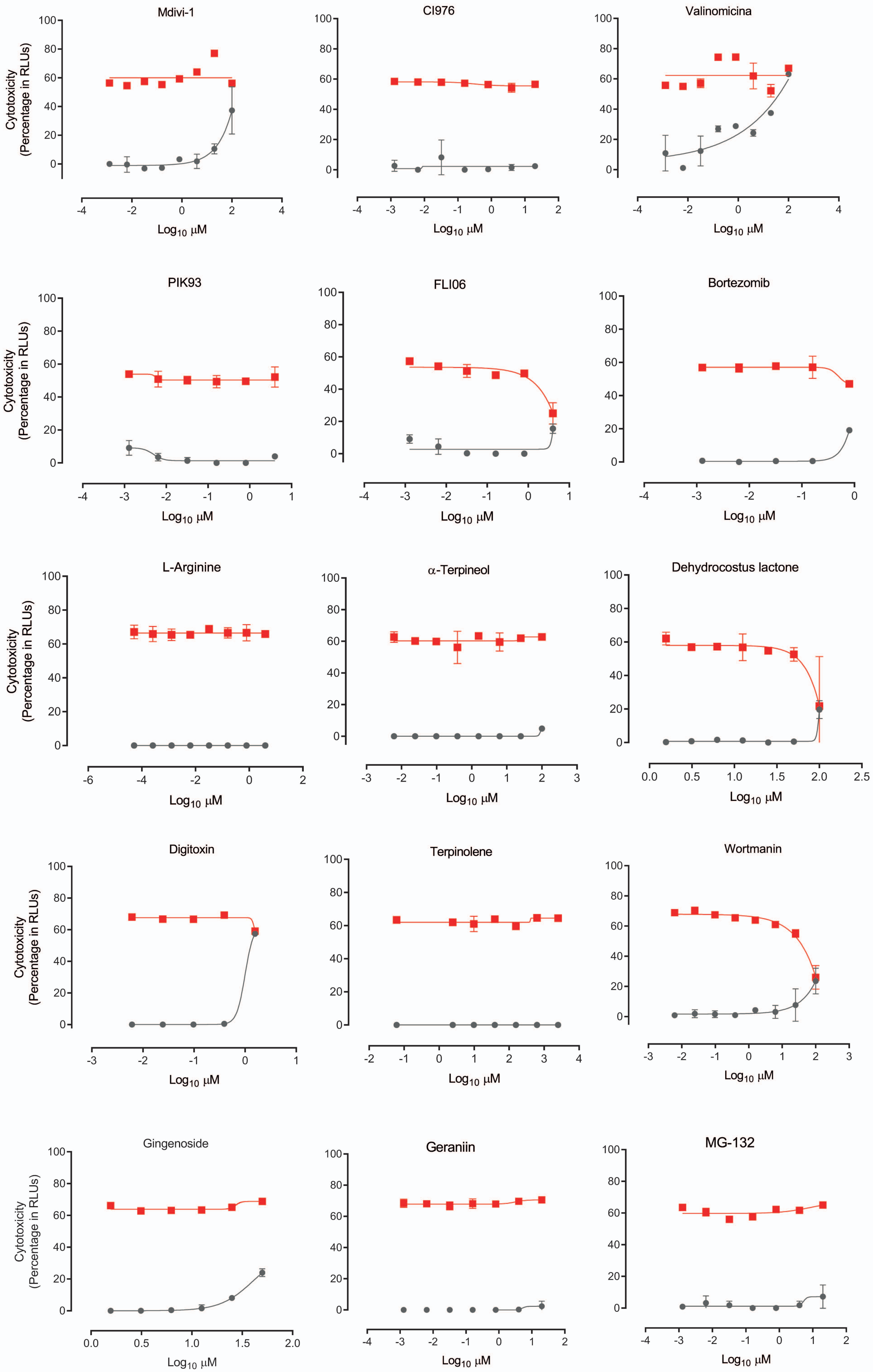

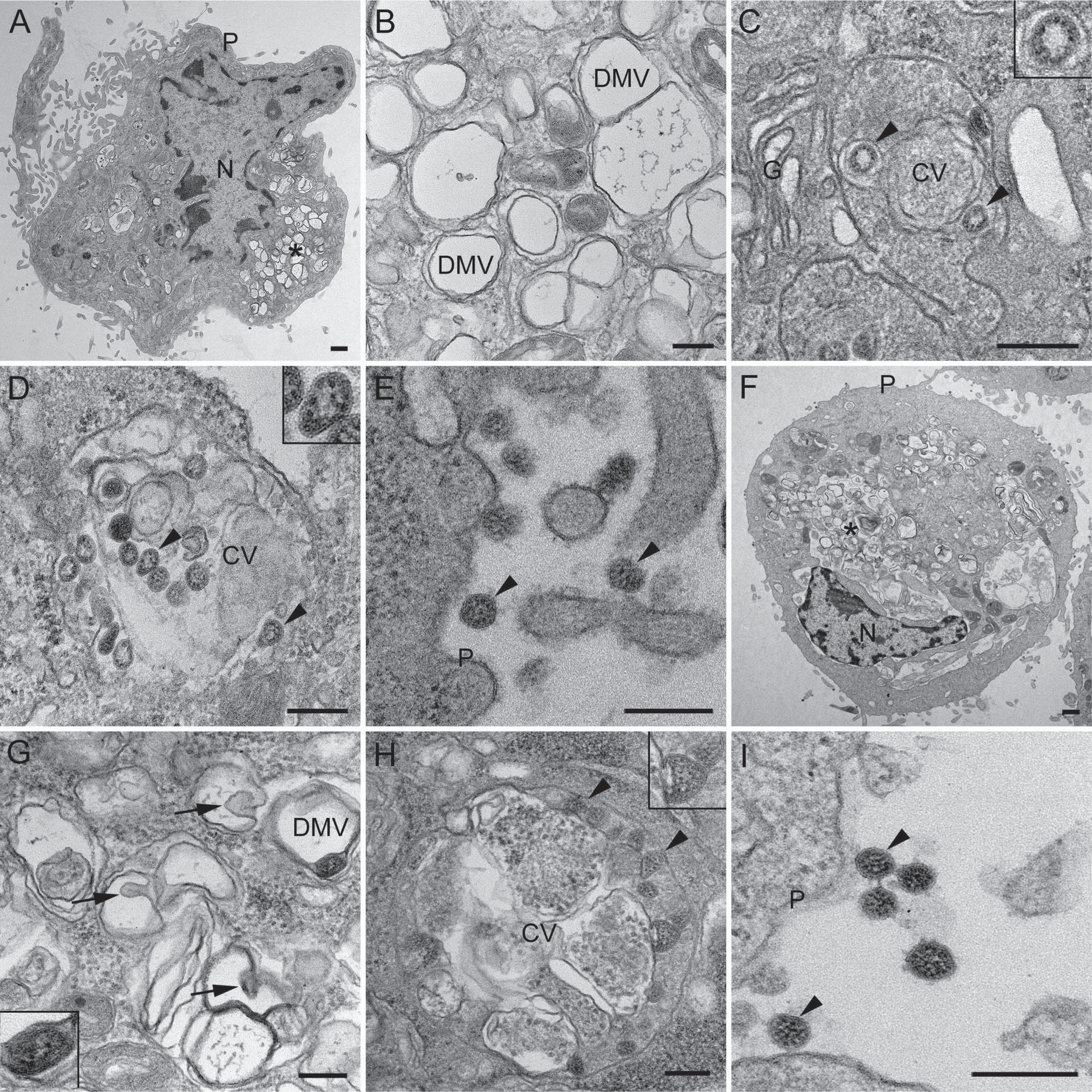
A
P
N
*
B
DMV
DMV
C
G
CV
D
CV
E
P
F
P
*
N
G
DMV
H
CV
I
P

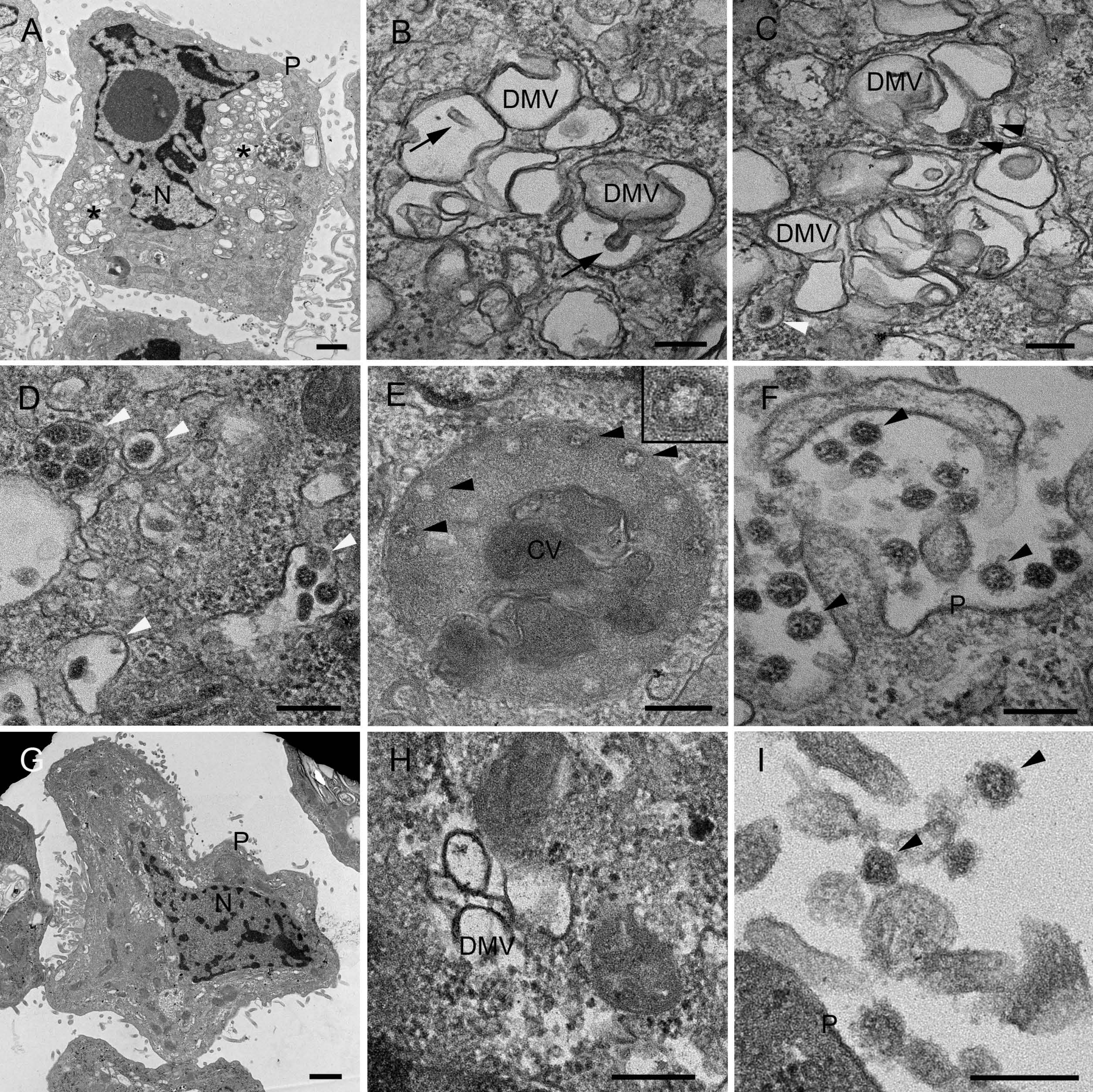

A
P
N
B
DMV
DMV
C
DMV
DMV
D
E
CV
F
P
G
P
N
H
DMV
I
P

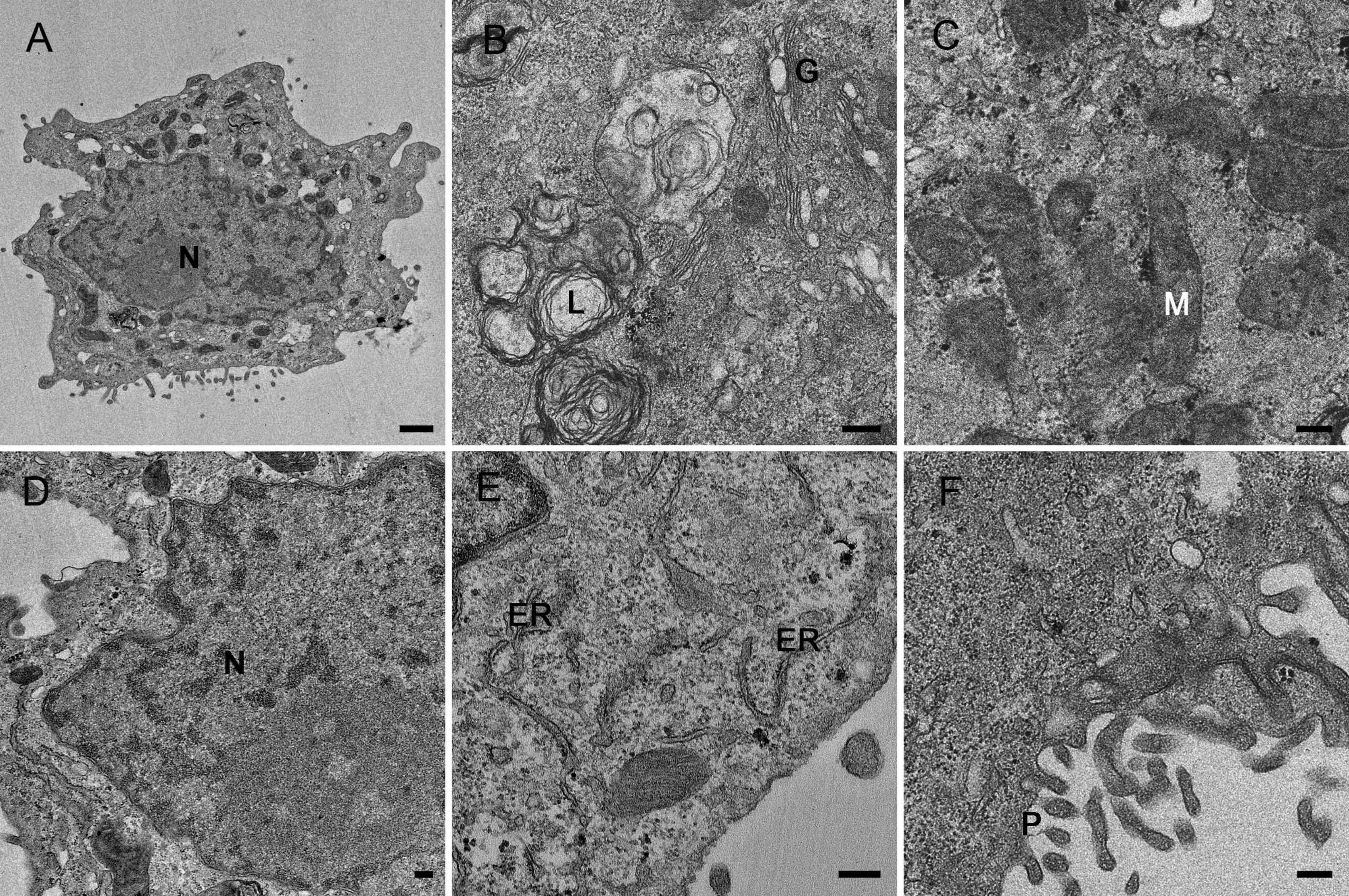
A
N
B
G
L
C
M
D
N
E
ER
ER
F
P

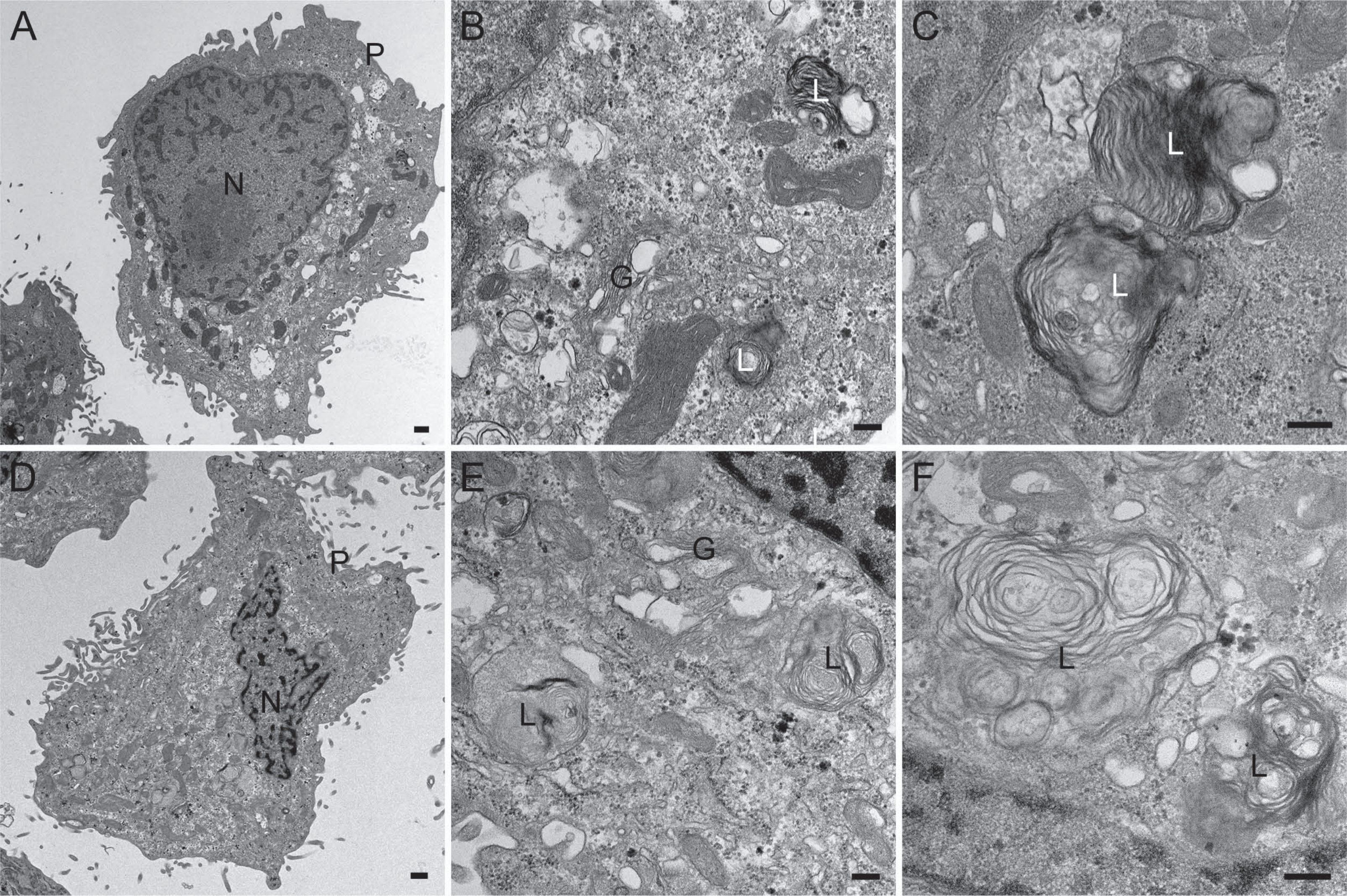
A
P
N
B
L
G
L
C
L
L
D
P
N
E
G
L
L
F
L
L

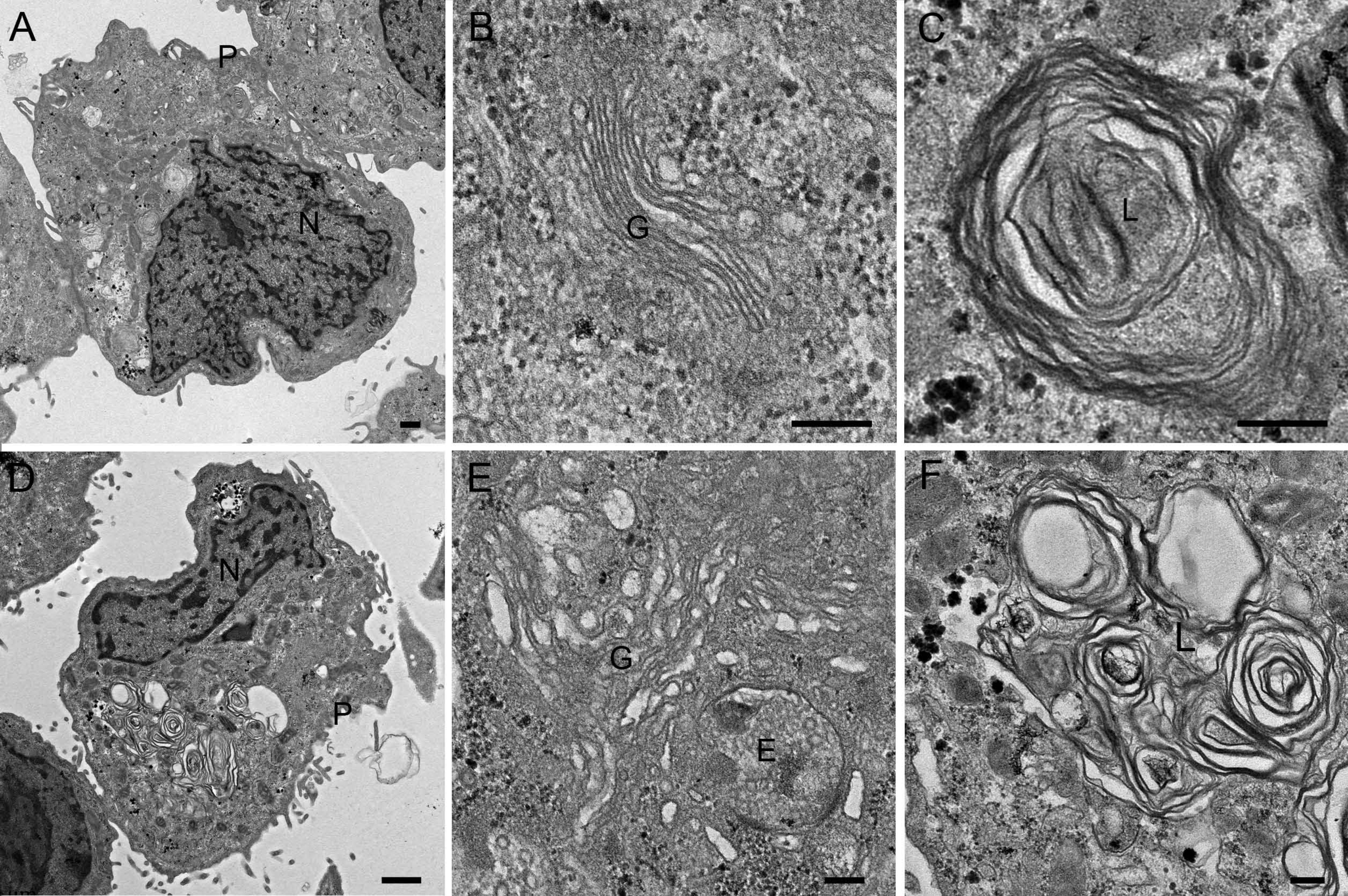
A
P
N
B
G
C
L
D
N
P
E
G
E
F
L

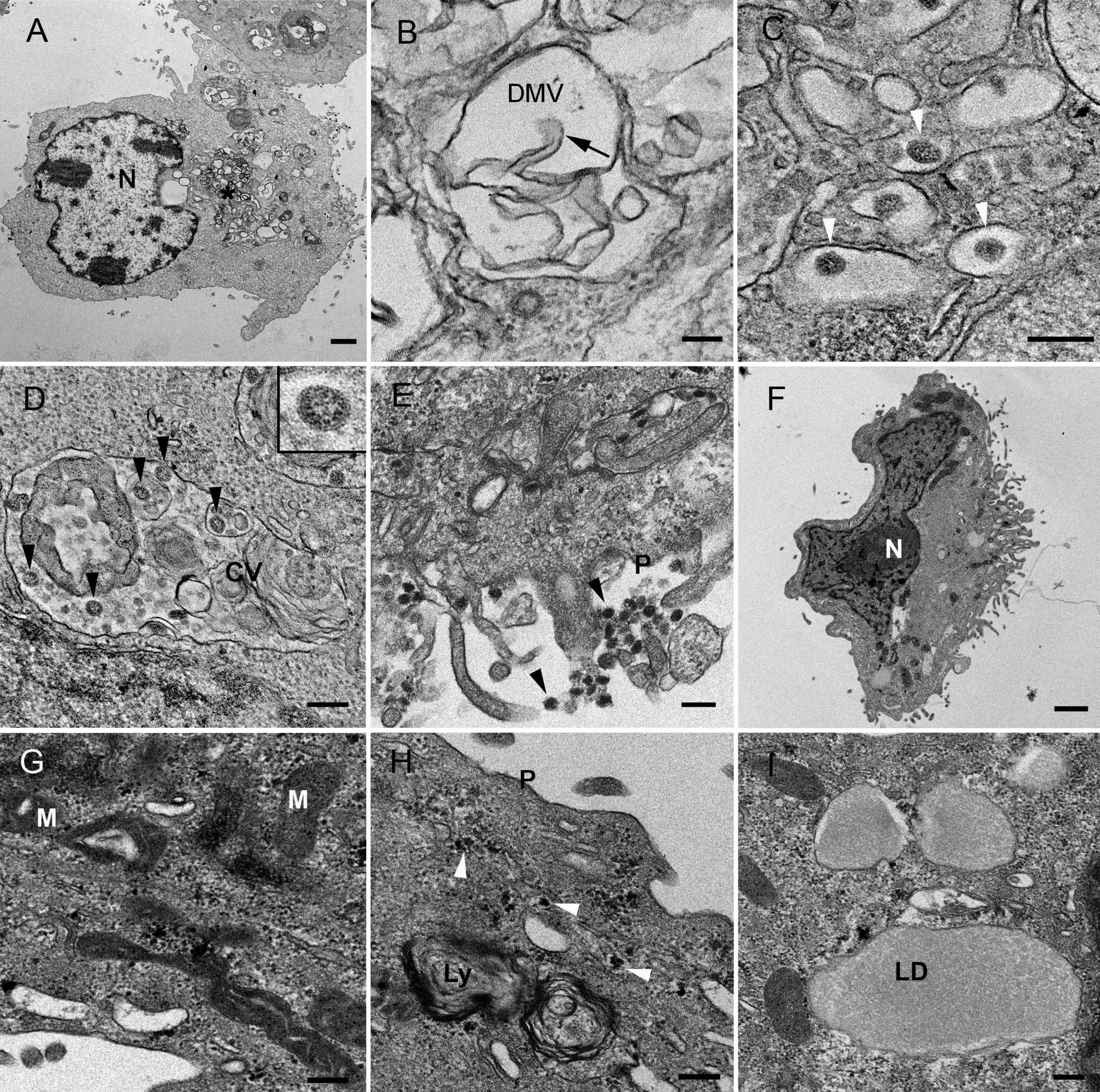

A
N
B
DMV
C
D
CV
E
P
F
N
G
M
M
H
P
Ly
I
LD

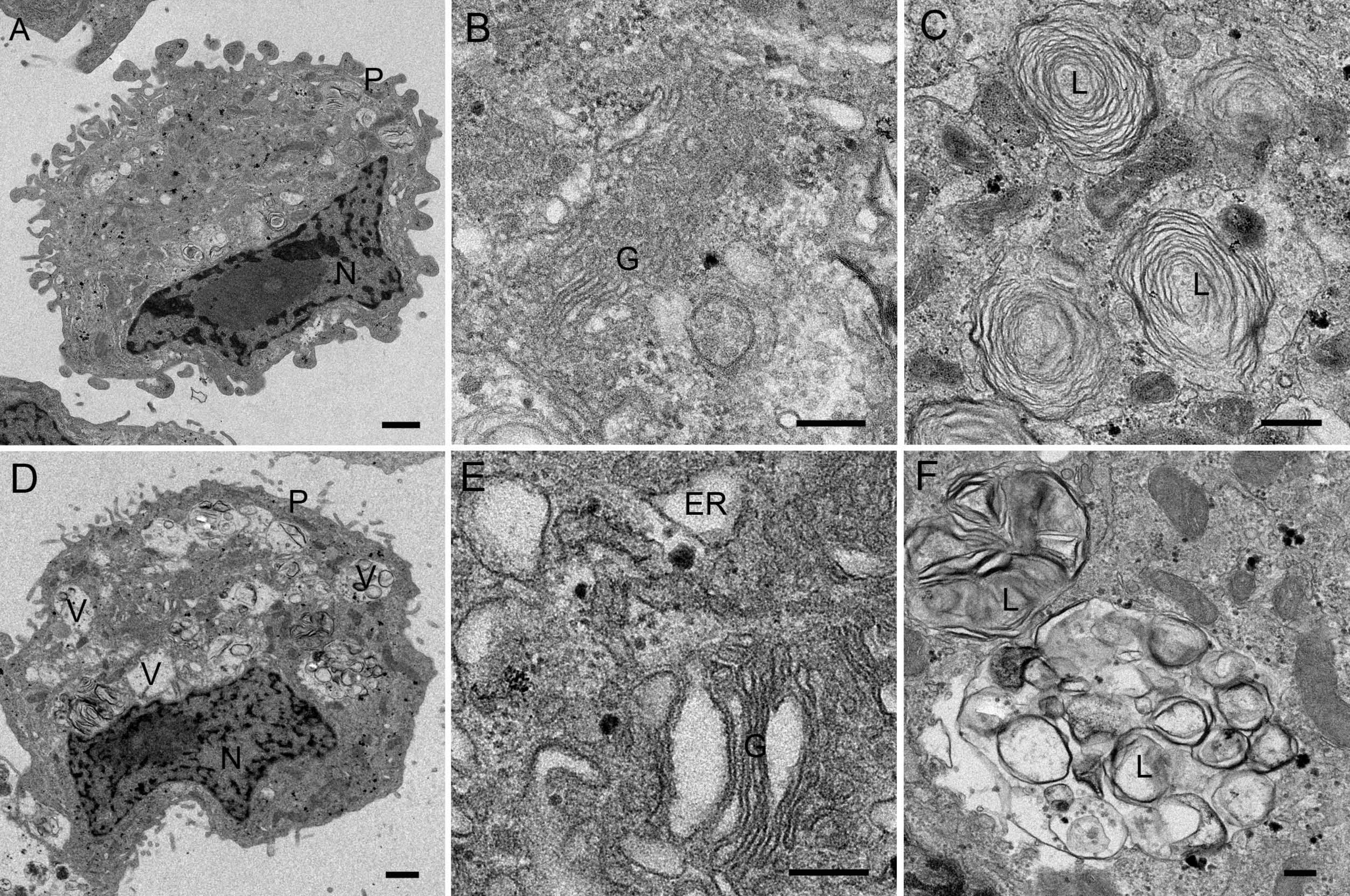
A
P
N
B
G
C
L
L
D
P
V
V
V
N
E
ER
G
F
L
L

### Supplementary Figure S9

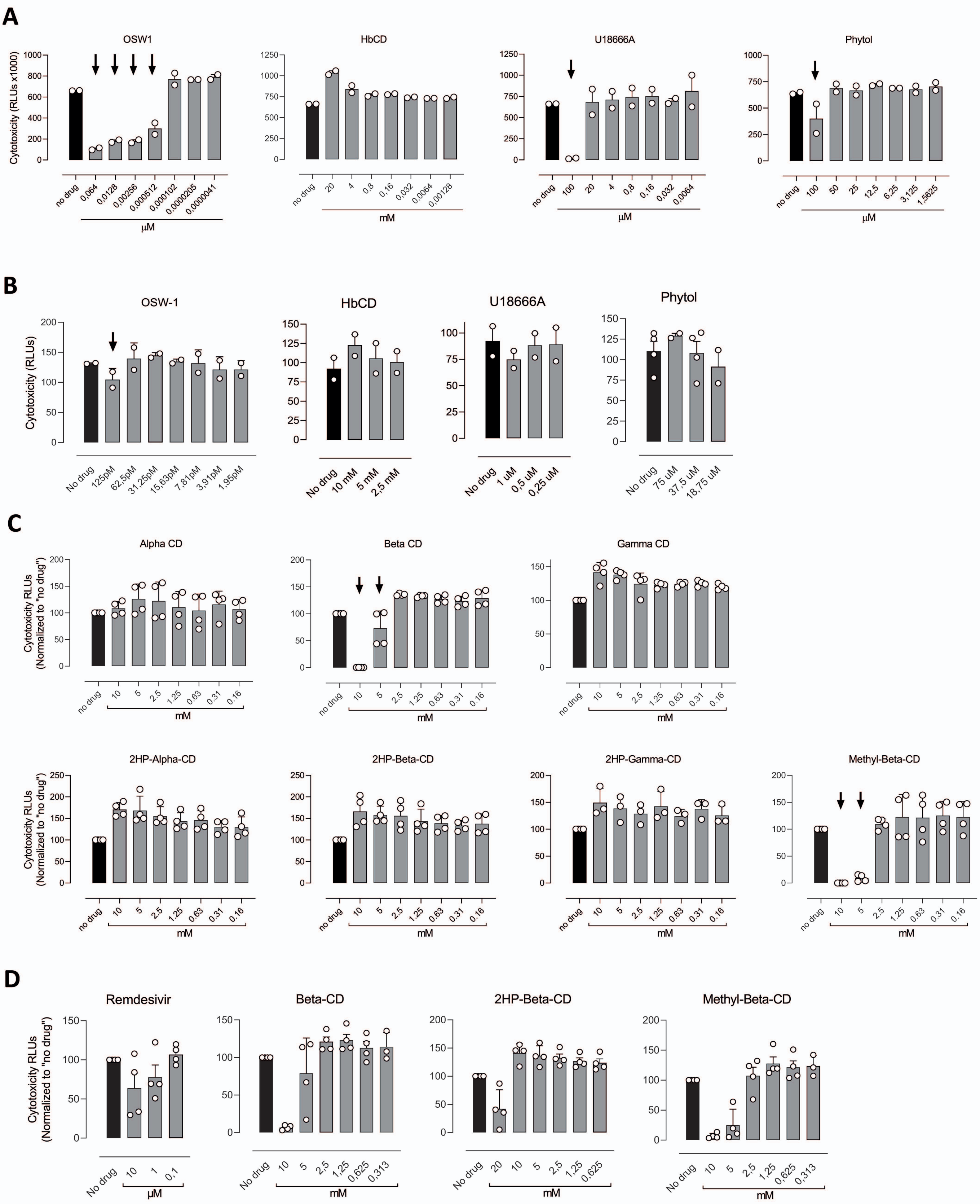

**SUPPLEMENTARY TABLES**

**Table S1.** Vendor origin and concentrations of tested cyclodextrins.

| Name | Vendor origin | Concentrations tested |
| --- | --- | --- |
| α-Cyclodextrin | Sigma-Aldrich (28705) | 10 – 0.16 mM |
| ß-Cyclodextrin | Sigma-Aldrich (W402826) | 10 – 0.16 mM |
| γ-Cyclodextrin | Sigma-Aldrich (C4892) | 10 – 0.16 mM |
| 2HP-α-Cyclodextrin | Sigma-Aldrich (390690) | 10 – 0.16 mM |
| 2HP-ß-Cyclodextrin | Sigma-Aldrich (H107) | 20 – 0.16 mM |
| 2HP-γ-Cyclodextrin | Sigma-Aldrich (H125) | 10 – 0.16 mM |
| Methyl-ß-Cyclodextrin | Sigma-Aldrich (C4555) | 10 – 0.16 mM |

**Table S2**. Energies for the top-ranked compounds against each indicated target.

| | Molecule | Binding Energy (kcal/mol) |
|---|---|---|
| Mpro | OSW1 | -108.79 |
| | Dispergo | -103.79 |
| | MG-132 | -99.44 |
| | Geraniin | -97.13 |
| | Proanthocyanidin-A2 | -89.11 |
| | Prodelphinidin-B3 | -83.42 |
| | PIK93 | -80.85 |
| PLPro | Grandinin* | -82.87 |
| | Silibin_A | -66.24 |
| | Itraconazole | -65.92 |
| | PIK93 | -65.06 |
| NPC1 | Itraconazole | -92.48 |
| | beta-Sitosterol | -88.98 |
| | U18666A | -85.69 |
| | Reserpine | -82.24 |
| | Arbidol | -80.62 |
| | Wortmannin | -77.72 |
| | Phytol | -77.27 |
| Spike | Rapamycin | -62.18 |
| | Dispergo | -61.82 |
| | OSW1 | -59.33 |
| | Curcumin | -58.67 |
| | Camostat mesylate | -58.51 |
| NSP16 | PIK93 | -50.04 |
| | Taxifolin | -47.83 |
| | Silibin A | -46.60 |
| | Myrcetin | -45.91 |
| | Quercetin | -44.64 |

(*) Grandinin does not fit into the pocket but interacts with the external surface of the target.

**Table S3.** Energies for the top-ranked cyclodextrins against the active site and the two allosteric sites of $M^{pro}$.

| | Molecule | Binding Energy (kcal/mol) |
|---|---|---|
| Active site | γ-CD | -91.87 |
| | HP-β-CD | -86.86 |
| | Methyl-β-CD | -86.63 |
| | β-CD | -81.77 |
| | α-CD | -65.47 |
| Allosteric site I | β-CD | -92.69 |
| | Methyl-β-CD | -78.76 |
| | γ-CD | -69.74 |
| | α-CD | -65.65 |
| Allosteric site II | HP-β-CD | -84.51 |
| | Methyl-β-CD | -83.61 |
| | γ-CD | -78.39 |
| | β-CD | -78.32 |
| | α-CD | -65.92 |

**Table S4.** Quantification of viral structures by TEM of Vero E6 cells treated with two doses (low and high) of each compound.

| | % cells with DMVs (% DMV positive cells with altered DMVs) | | % of cells with SMVs (#/cell) | | % of cells with CV (#/cell) | | % of cells with extracellular VPs (#/cell) | |
|---|---|---|---|---|---|---|---|---|
| **Control** | 75% (40%) | | 80% (3.6/cell) | | 40% (0.7/cell) | | 100% (40.9/cell) | |
| | low | high | low | high | low | high | low | high |
| **OSW 1** | 85% (18%) | 65% (92%) | 90% (3.3/cell) | 60% (2.8/cell) | 75% (2.5/cell) | 75% (2.9/cell) | 100% (34.4/cell) | 80% (23.8/cell) |
| **HβCD** | 32% (86%) | 22% (20%) | 82% (3.5/cell) | 26% (1.4/cell) | 45% (1.3/cell) | 13% (0.4/cell) | 95% (34/cell) | 65% (8/cell) |
| **U18666A** | 85% (82%) | 36%* (100%) | 80% (4.8/cell) | 0% (0/cell) | 55% (1/cell) | 0% (0/cell) | 100% (42/cell) | 14% (0.2/cell) |
| **Phytol** | 60% (42%) | x | 60% (2.8/cell) | x | 20% (0.5/cell) | x | 100% (58.85/cell) | x |
